# *De novo* DNA methylation controls neuronal maturation during adult hippocampal neurogenesis

**DOI:** 10.1101/2020.09.22.308692

**Authors:** Sara Zocher, Rupert W Overall, Gabriel Berdugo-Vega, Nicole Rund, Anne Karasinsky, Vijay S Adusumilli, Christina Steinhauer, Sina Scheibenstock, Kristian Händler, Joachim L Schultze, Federico Calegari, Gerd Kempermann

**Affiliations:** German Center for Neurodegenerative Diseases (DZNE) Dresden, Tatzberg 41, 01307 Dresden, Germany; Center for Regenerative Therapies Dresden (CRTD), Technische Universität Dresden, Fetscherstraße 105, 01307 Dresden, Germany; PRECISE Platform for Single Cell Genomics and Epigenomics, German Center for Neurodegenerative Diseases and University of Bonn, 53175 Bonn, Germany

**Keywords:** Neural stem cells, adult neurogenesis, neuron function, synaptogenesis, DNA methylation, DNA methyltransferases, Dnmt3a, epigenetic, hippocampus, environmental enrichment

## Abstract

Dynamic DNA methylation controls gene-regulatory networks underlying cell fate specification. How DNA methylation patterns change during adult hippocampal neurogenesis and their relevance for adult neural stem cell differentiation and related brain function has, however, remained unknown. Here, we show that neurogenesis-associated *de novo* DNA methylation is critical for maturation and functional integration of adult-born hippocampal neurons. Cell stage-specific bisulfite sequencing revealed a pronounced gain of DNA methylation at neuronal enhancers, gene bodies and binding sites of pro-neuronal transcription factors during adult neurogenesis, which mostly correlated with transcriptional up-regulation of the associated loci. Inducible deletion of both *de novo* DNA methyltransferases *Dnmt3a* and *Dnmt3b* in adult neural stem cells specifically impaired dendritic outgrowth and synaptogenesis of new-born neurons, resulting in reduced hippocampal excitability and specific deficits in hippocampus-dependent learning and memory. Our results highlight that, during adult neurogenesis, remodeling of neuronal methylomes is fundamental for proper hippocampal function.

## Introduction

The dentate gyrus of the hippocampus harbors a stem cell population that generates new neurons throughout life (Bond et al., 2015; Kempermann et al., 2015). Adult-born neurons are critical for hippocampus-dependent cognitive flexibility and impairments in their production have been implicated in age-related cognitive decline and neuropsychiatric disorders such as depression (Anacker et al., 2018; Berdugo-Vega et al., 2020; Toda et al., 2019). During adult neurogenesis, cell-intrinsic molecular programs and environmental signals interact to control neural stem cell proliferation, neuronal differentiation, maturation and functional integration of the new neurons into the hippocampal circuitry (Aimone et al., 2014; Vicidomini et al., 2020). Adult neural stem cell differentiation is associated with transient and irreversible gene expression changes which are crucial for neurogenic commitment and maintenance of the neuronal identity (Vicidomini et al., 2020). Although a number of underlying transcription factors have been identified (Beckervordersandforth et al., 2015; Mukherjee et al., 2016; Schäffner et al., 2018), the epigenetic machinery that integrates cell-intrinsic and extrinsic signals to enable the molecular changes required for the generation of functional neurons in the adult hippocampus has remained elusive.

DNA methylation is an epigenetic modification with established roles in stem cell differentiation and lineage commitment (Mohn and Schübeler, 2009; Smith and Meissner, 2013). For instance, DNA methylation is involved in the long-term silencing of multipotency and proliferation genes, the control of lineage-specific transcription factors and the maintenance of new cell type identities (Beerman and Rossi, 2015; Challen et al., 2012; Smith and Meissner, 2013). The genomic regions that change DNA methylation during adult hippocampal neurogenesis are unknown, but - given the importance of DNA methylation in creating neuronal diversity (Mo et al., 2015) - promise prominent insight into the gene-regulatory networks underlying this exceptional form of brain plasticity. A functional role of DNA methylation changes during adult neurogenesis has previously been suggested by the disruption of the process through knock-out of the demethylating enzymes Tet1 and Tet2 as well as the methylation readers Mbd1 and Mecp2 (Gontier et al., 2018; Jobe et al., 2017; Li et al., 2014; Zhang et al., 2013). Moreover, both the neurogenic activity of adult hippocampal stem cells as well as brain DNA methylation patterns are sensitive to environmental stimuli and behavioral activity (Van Praag et al., 2000; Zocher et al., 2020). This makes DNA methylation an exciting potential mechanism underlying experience-dependent brain plasticity.

DNA methylation is catalyzed by the activity of DNA methyltransferases. Whereas Dnmt1 maintains global DNA methylation patterns during replication, *de novo* DNA methyltransferases Dnmt3a and Dnmt3b catalyze the addition of new methyl groups to previously non-methylated genomic regions (Ma et al., 2010). *De novo* DNA methyltransferases are crucial for embryonic and postnatal brain development and memory formation in adulthood (Gulmez Karaca et al., 2020; Odell et al., 2020; Wu et al., 2010), but their specific role during adult neurogenesis has been as yet unknown. Previous studies suggested that, in neural precursor cells derived from *Dnmt3a*-deficient embryonic stem cells, astrogliogenesis was increased at the expense of neurogenesis (Wu et al., 2010, 2012; Ziller et al., 2018). However, due to the lack of inducible *Dnmt3a* knock-out models it remained unknown whether a neurogenic fate of neural precursor cells is pre-defined by DNA methylation patterns established during development or whether it is specified by further *de novo* DNA methylation in the course of terminal neuronal differentiation.

Here, we provide a comprehensive molecular, cellular and functional characterization of the DNA methylation changes during *in vivo* adult hippocampal neurogenesis, their sensitivity to environmental stimulation and their relevance for behavior and cognition. By generating an inducible, neural stem cell-specific knock-out model of *de novo* DNA methyltransferases, we demonstrate the specific requirement of neurogenesis-associated *de novo* DNA methylation for the maturation and functional integration of adult-born neurons into the hippocampus.

## Results

### Prominent *de novo* DNA methylation of neuronal genes during adult neurogenesis

To characterize the DNA methylation changes associated with adult neurogenesis, we isolated Nestin-expressing neural stem and early progenitor cells (NSPCs), Doublecortin (Dcx)- expressing late progenitor cells (lPCs) and NeuO-positive neurons from the dentate gyrus of adult mice using fluorescence activated cell sorting (FACS; Fig. 1A; Fig. S1A-E). To further define cell types from the heterogenous pool of Dcx-positive cells, we established a protocol to specifically isolate progenitor cells from neurons by making use of their distinct scattering properties (Fig. S1F-G). We confirmed the neurogenic trajectory of our isolated cell populations by quantitative polymerase chain reaction, which showed an expected successive down-regulation of proliferation and precursor cell markers from NSPCs to neurons (Fig. S1H).

**Figure 1:**
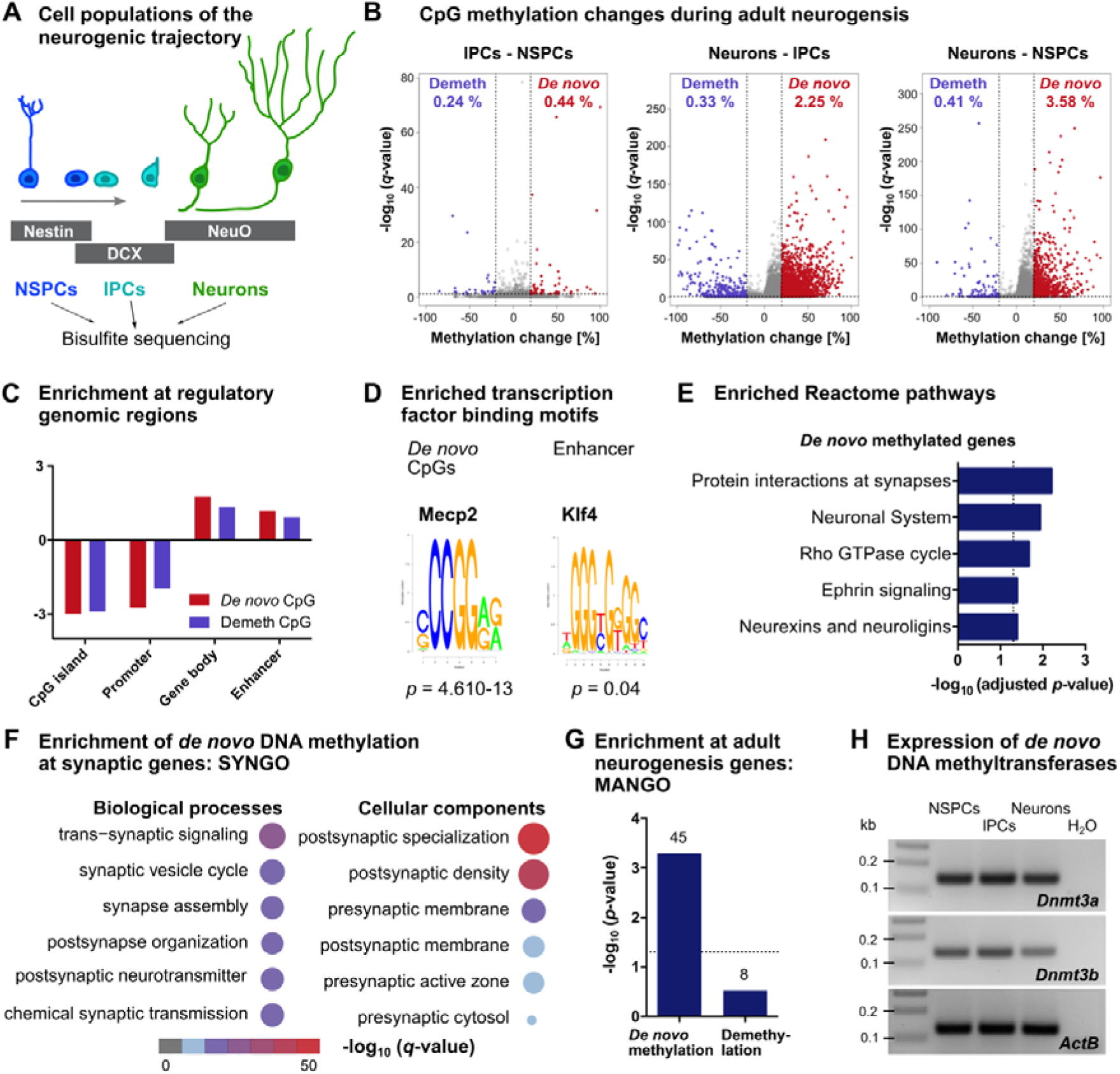
DNA methylation changes during adult hippocampal neurogenesis. **A,** Neural stem and early progenitor cells (NSPCs), late neural progenitor cells (lPCs) and neurons were isolated from the dentate gyrus of 10-week-old mice using the markers Nestin, Doublecortin (DCX) and NeuO, respectively. DNA methylation profiles of each cell population were generated by Reduced Representation Bisulfite Sequencing (RRBS; n = 5 – NSPCs; 6 – lPCs; 4 – neurons). **B,** Volcano plots depicting CpG methylation changes between cell populations. Significantly differentially methylated cytosines with *q* < 0.05 and absolute methylation differences greater than 20 % are highlighted in violet (demethylated CpGs – Demeth) and red (*de novo* methylated CpGs - *De novo*). Percentages of differentially methylated CpGs among all sequenced CpGs are indicated with the respective color. **C,** Neurogenesis-associated differentially methylated CpGs are depleted at promoters and CpG islands but enriched at gene bodies and enhancers. Adjusted *p* < 0.001 for all bars. **D,** Top enriched transcription factors from transcription factor motif enrichment with all *de novo* methylated CpGs (3,151 CpGs) and *de novo* methylated CpGs located within enhancer regions (561 CpGs). Depicted *p*-values are adjusted *p*-values from hypergeometric tests. **E,** Significantly enriched pathways of *de novo* methylated genes (2,048 genes) compared to all sequenced genes. **F,** Top 5 enriched biological processes (left) and cellular components (right) of *de novo* methylated genes from gene set enrichment analysis using the Synaptic Gene Ontologies (SYNGO). Size of dots represents log_2_ (number of genes in term). **G,** Genes with known function in adult hippocampal neurogenesis as annotated in the Mammalian Adult Neurogenesis Gene Ontology (MANGO) are significantly enriched among *de novo* methylated genes. Numbers of overlapping genes are indicated on top of the respective bar. **H,** Transcript expression of *de novo* DNA methyltransferases *Dnmt3a* and *Dnmt3b* in neural precursor cells and neurons as determined by RT-PCR.

To identify DNA methylation changes during neuronal differentiation, we performed Reduced Representation Bisulfite Sequencing (RRBS; Boyle et al., 2012) and obtained single-nucleotide resolution DNA methylation profiles of NSPCs, lPCs and neurons. Comparing CpG methylation levels between the cell populations, we found that, while only 0.68 % of CpGs changed methylation from NSPCs to lPCs, 2.58 % of CpGs were differentially methylated between lPCs and neurons (Fig. 1B). In total, we identified 3,621 CpGs that significantly changed methylation during neuronal differentiation in the hippocampus (NSPCs vs lPCs or lPCs vs neurons or NSPCs vs neurons; Supplemental data 1). Among these CpGs, the majority (87.0 %) gained methylation (*de novo* methylated CpGs) during neuronal differentiation. Both *de novo* methylated and demethylated CpGs were depleted at promoters and CpG islands, but significantly enriched at gene bodies and enhancers (Fig. 1C). Most *de novo* methylated enhancers (84.01 %) were located within gene bodies, compared to only 15.99 % that were found at distal intergenic regions. To investigate a potential gene-regulatory role of the detected methylation changes, we performed transcription factor motif enrichment. The strongest enriched motif among *de novo* methylated CpGs corresponded to the methyl-CpG-binding protein Mecp2 (Fig. 1D), which binds methylated cytosines broadly across the genome and is known to control adult neurogenesis (Li et al., 2014). When we restricted the analysis to *de novo* methylated CpGs located within enhancers, the only enriched transcription factor motif was an alternative, methylation-dependent binding site of Klf4 (Hu et al., 2013). Binding of Klf4 to the methylated sequence has been shown to mediate transcriptional activation related to cell migration and morphogenesis (Wan et al., 2017). These results suggested prominent *de novo* DNA methylation of gene-regulatory regions during adult neurogenesis, including binding sites of pro-neuronal transcription factors such as Mecp2 and Klf4.

Although we only detected 113 CpHs (cytosines followed by any other nucleobase than guanine) which significantly changed methylation during neuronal differentiation (Fig. S2A-B), neurons showed higher global CpH methylation levels compared to NSPCs and lPCs (Fig. S2C). This observation is consistent with the previously reported accumulation of CpH methylation during neuronal differentiation (Mohn et al., 2008). We also observed an increase in global CpG methylation during neuronal differentiation (Fig. S2D), supporting the predominant *de novo* methylation found at the individual CpG level.

To gain first insight into the functional role of *de novo* DNA methylation, we annotated differentially methylated CpGs to the nearest located gene and performed a Reactome pathway analysis with the 2,024 genes containing *de novo* methylated CpGs. *De novo* methylated genes were significantly enriched at pathways involved in neuronal maturation and synaptic plasticity, such as RhoGTPase cycle, ephrin signaling and neurexins (Fig. 1E). To confirm the enrichment in synapse-related pathways, we used the Synaptic Gene Ontologies (SYNGO) – an expert-curated knowledgebase of synaptic genes (Koopmans et al., 2019). In total, 148 *de novo* methylated genes were annotated in SYNGO with enrichment in sub-ontologies related to synaptogenesis and synaptic function (Fig. 1F). Additional gene set enrichment analyses showed that *de novo* methylated loci were also significantly enriched at genes with known function in adult hippocampal neurogenesis as annotated in the Mammalian Adult Neurogenesis Gene Ontology (MANGO; Overall et al., 2012; Fig. 1G). Together, these results indicated that neuronal differentiation during adult hippocampal neurogenesis is associated with *de novo* DNA methylation of genes with known relevance in neuronal maturation and synaptic plasticity.

### Deletion of *de novo* DNA methyltransferases influences neuronal differentiation *in vitro*

We next sought to dissect the functional role of neurogenesis-associated *de novo* DNA methylation through the deletion of *de novo* DNA methyltransferases in NSPCs. First, we confirmed that *Dnmt3a* and *Dnmt3b* are expressed in NSPCs, lPCs and neurons of the adult hippocampus (Fig. 1H). We then generated a mouse model that enables the conditional and inducible deletion of both *de novo* DNA methyltransferases during adult hippocampal neurogenesis by crossing nestinCre-ER^T2^ mice (Imayoshi et al., 2013) with Dnmt3a^(fl/fl)^ (Kaneda et al., 2004) and Dnmt3b^(fl/fl)^ mice (Dodge et al., 2005) (Fig. 2A). In this mouse model, administration of tamoxifen during adulthood would lead to gene knock-out in adult NSPCs and their differentiated progeny, but importantly, would not affect brain development.

**Figure 2:**
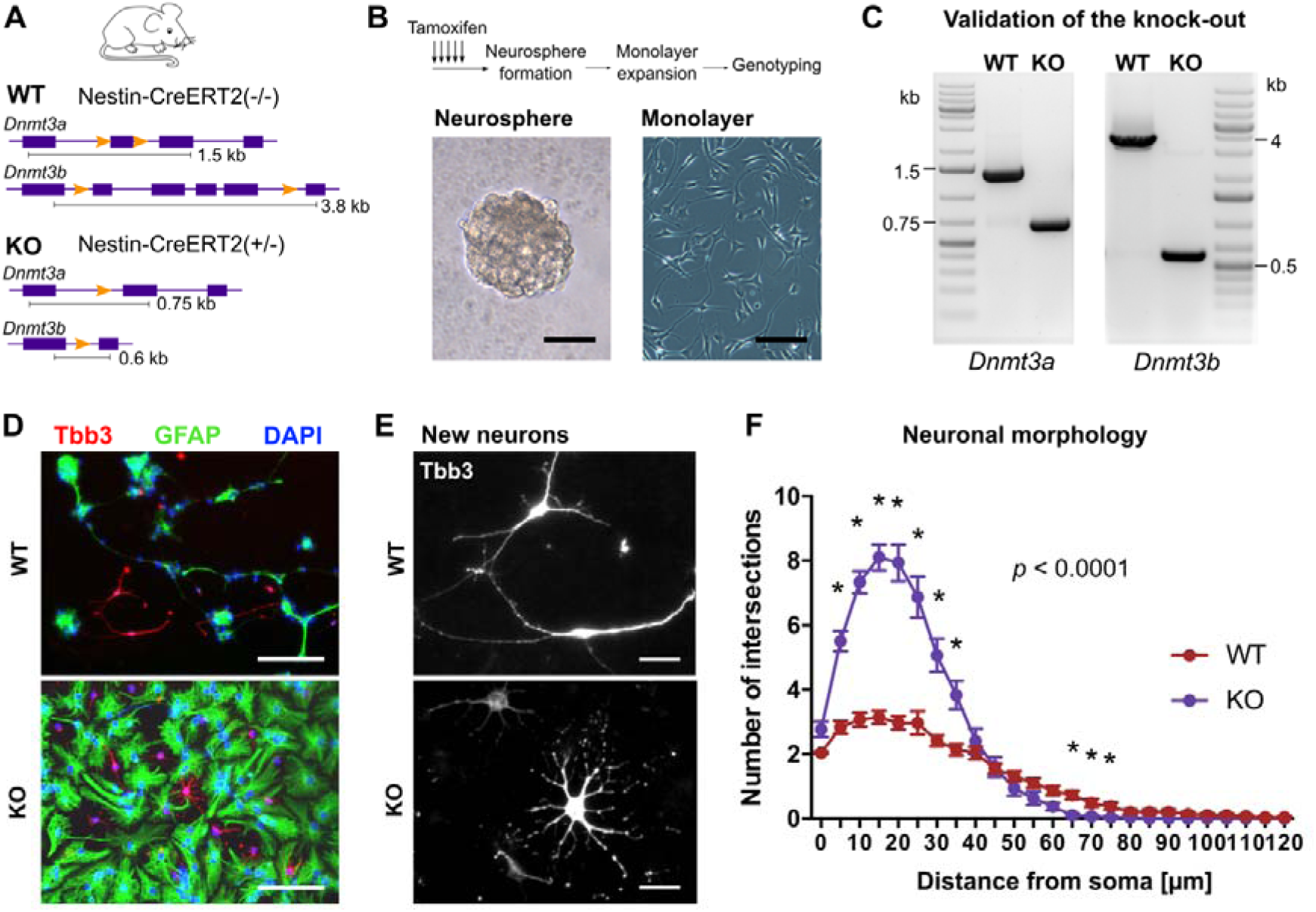
Deletion of *de novo* DNA methyltransferases in neural precursor cells influences neuronal maturation *in vitro*. **A,** Schematic representation of the genomic area of *Dnmt3a* and *Dnmt3b* in wildtype mice (WT) and knock-out mice (KO). WT were *Dnmt3a*^fl/fl^ / *Dnmt3b* ^fl/fl^ / nestin-C*reERT2* (-/-), while KO were *Dnmt3a* ^fl/fl^ / *Dnmt3b* ^fl/fl^ / nestin-CreER^T2^ (+/-). **B,** At an age of six weeks, WT and KO received five injections of tamoxifen over a period of five days. To generate clonal neural precursor cell lines, dentate gyri were dissociated 10 days after the first tamoxifen injection and plated for neurosphere formation. Monolayer cultures were generated by dissociating single neurospheres. Representative pictures of neurosphere (left) and neurosphere-derived adherent monolayer culture (right). Scale bars: 50 µm. **C,** Representative images of polymerase chain reaction used for genotyping of clonal neural precursor cell lines. **D,** Neural precursor cells from WT and KO mice are able to differentiate into astrocytes (Gfap-positive) and neurons (Tbb3-positive). Scale bar: 100 µm. **E,** High magnification fluorescent image of differentiated neurons in WT and KO cultures. Scale bar: 20 µm. **F,** Sholl analysis of Tbb3-labeled WT and KO neurons (n = 29, WT; n = 30, KO). Reported *p*–value corresponds to genotype effect from two-way ANOVA. Asterisks highlight data points with *p* < 0.05 after multiple testing adjustment of repeated *t*-tests. Depicted are means ± standard errors of the mean.

Previous studies analyzed the role of *de novo* DNA methyltransferases in *Dnmt3a*-deficient embryonic stem cells during neuronal differentiation in culture (Wu et al., 2010, 2012; Ziller et al., 2018). To compare our results with these studies, we first investigated whether inducible deletion of *de novo* DNA methyltransferases in NSPCs influenced their proliferation and differentiation potential *in vitro*. We therefore administered tamoxifen to wildtype (WT) and knock-out (KO) mice and generated neural precursor cell cultures from their dentate gyri (Fig. 2B). To obtain cultures with homozygous KO of *Dnmt3a* and *Dnmt3b*, we exploited the ability of single NSPCs to grow as neurospheres (Walker et al., 2016) and expanded individual neurospheres to obtain clonal monolayer cultures. To characterize the knock-out efficiency of our nestinCre-ER^T2^::Dnmt3a^(fl/fl)^-Dnmt3b^(fl/fl)^ mouse line, we genotyped the clonal cultures using polymerase chain reaction (Fig. 2C). We estimated that on average 82.35 % ± 4.79 of activatable (neurosphere-forming) NSPCs in KO mice contained homozygous deletions of both *Dnmt3a* and *Dnmt3b* (mean ± standard error of the mean; n = 5 mice).

No difference in proliferation was observed between WT and KO cultures (Fig. S3A-C), suggesting that *de novo* DNA methyltransferases are dispensable for *in vitro* neural precursor cell proliferation. Neural precursor cells from WT and KO mice both differentiated into neurons and astrocytes after withdrawal of epidermal growth factor (EGF) and fibroblast growth factor (FGF-2) from the culture medium (Fig. 2D). However, differentiated neurons from KO cultures showed clear morphological differences compared to WT neurons (Fig. 2E), which we confirmed by Sholl analysis (Fig. 2F). While WT neurons developed 2-4 elongated processes, neurons from KO cultures exhibited 8-12 processes with shorter dendritic length. Together, these results showed that *de novo* DNA methyltransferases influence the maturation of newly-formed neurons *in vitro*.

### *De novo* DNA methyltransferases establish neuronal DNA methylation patterns during neurogenesis

To investigate the extent of Dnmt3a/b-dependent DNA methylation during neuronal differentiation, we isolated *in vitro* differentiated neurons by FACS and compared DNA methylation profiles between WT and KO neurons using RRBS (Fig. 3A). We identified focal hypomethylation at 21,109 CpGs in KO compared to WT neurons (Fig. 3B; Supplemental data 2), identifying those cytosines as targets of Dnmt3a and/or Dnmt3b. Hypomethylated CpGs in KO neurons were significantly enriched at gene bodies and enhancers (Fig. 3C). In total, Dnmt3a/b-dependent hypomethylation targeted 1,554 enhancers and 154 neuronal super-enhancers (Fig. 3D), indicating that *de novo* DNA methyltransferases mediate *de novo* DNA methylation of enhancers during neurogenesis.

**Figure 3:**
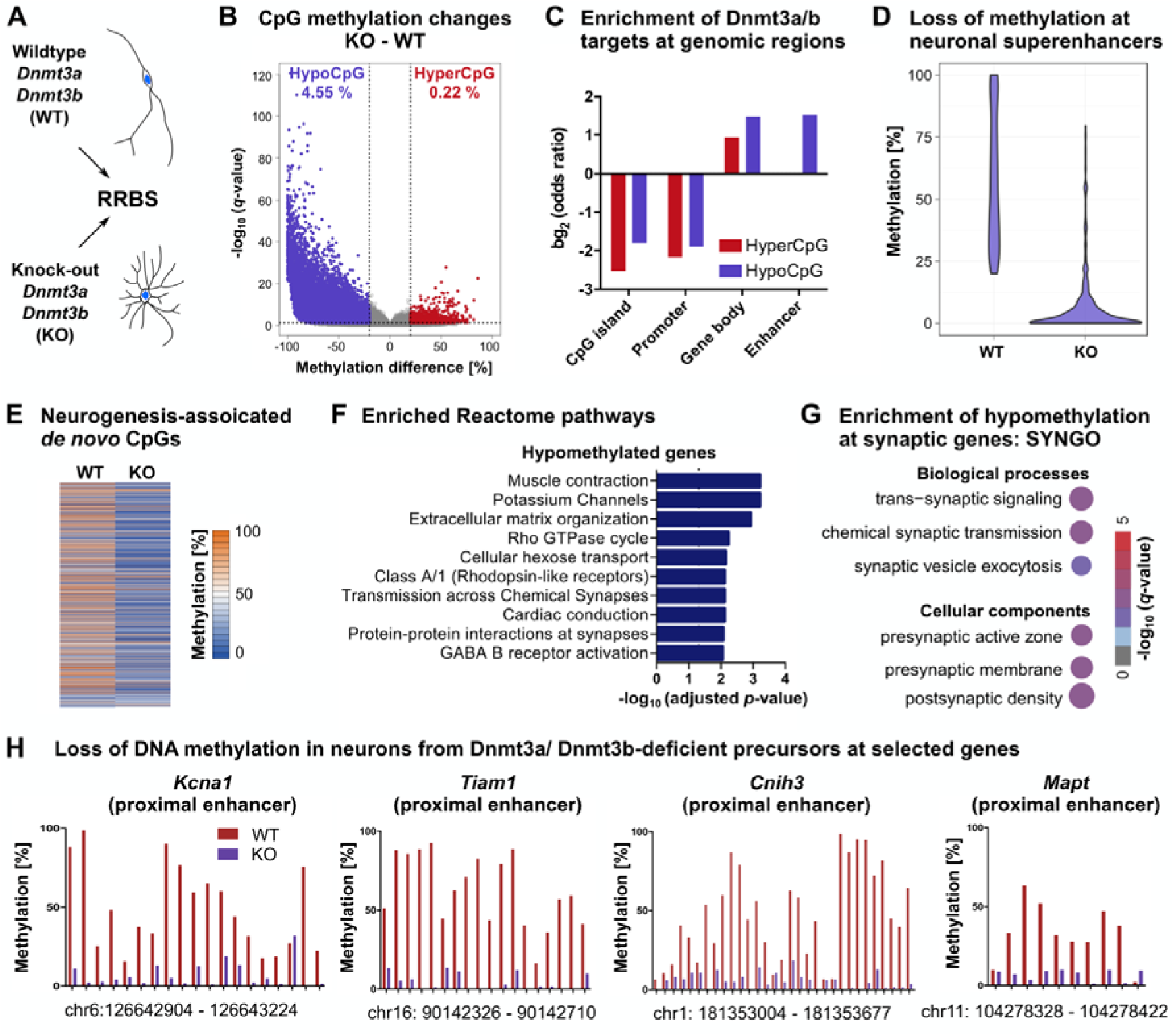
*De novo* DNA methyltransferases establish neuronal DNA methylation patterns. **A,** Neural precursor cells with and without knock-out of *Dnmt3a* and *Dnmt3b* were differentiated into neurons. Neurons were isolated by FACS and the DNA methylation patterns compared between WT and KO using RRBS (n = 3). **B,** In total, 22,150 CpGs significantly changed methylation (*q* < 0.05 and absolute methylation difference > 20 %) between KO and WT neurons, accounting to 4.55 % of all CpGs being hypomethylated (hypoCpG; violet) and 0.22 % being hypermethylated (hyperCpG; red) in KO neurons. **C,** Hyper- and hypomethylated CpGs were depleted from CpG islands and promoters, but enriched at gene bodies. Hypomethylated but not hypermethylated CpGs were enriched at enhancer regions (adjusted *p* < 0.001). **D,** Hypomethylated CpGs were also enriched at neuronal super-enhancers (*p* < 0.001). Depicted are methylation percentages of the 615 hypomethylated CpGs located within neuronal super-enhancers. **E,** CpGs that gained methylation during adult neurogenesis *in vivo* (see Fig. 1), were hypomethylated in neurons from KO compared to WT cultures. **F,** Genes containing hyomethylated cytosines (6,275 genes) were significantly enriched in pathways related to neuronal function. **G,** Significantly enriched biological processes and cellular components from SYNGO enrichment analysis. **H,** Loss of DNA methylation at representative neuronal genes in KO neurons. Depicted genomic regions are specified underneath the plots. Bars represent means.

In addition to hypomethylated CpGs, we found 1,041 CpGs that gained methylation through KO of *de novo* DNA methyltransferases (4.70 % of differentially methylated CpGs) that were enriched at gene bodies but not at enhancers. Isolated hypermethylation after deletion of DNA methyltransferases has been reported before (Charlton et al., 2020) and likely represents an indirect consequence of deleting *de novo* DNA methyltransferase activity through their interaction with other molecular players.

CpGs that gained methylation during neuronal differentiation *in vivo* significantly overlapped with hypomethylated CpGs in KO neurons (hypergeometric test: *p* < 0.001) and showed a general loss of methylation in KO compared to WT neurons (Fig. 3E; Fig. S4). This confirmed that neurogenesis-associated *de novo* DNA methylation is mediated by the canonical *de novo* DNA methyltransferases Dnmt3a and/ or Dnmt3b. Deletion of Dnmt3a/b induced hypomethylation at 6,275 genes, which were enriched in pathways involved in neuronal function, including potassium channels, Rho GTPase cycle and synapse-related pathways (Fig. 3F), and at SYNGO terms related to synaptic signaling and synaptogenesis (Fig. 3G). These included enhancers of core neuronal genes, such as *Tiam1* and *Mapt*, which are both involved in neuronal growth and axon guidance, as well as the potassium channel encoding gene *Kcna1* and the glutamate receptor modulator *Cnih3* (Fig. 3H).

These results indicated that *de novo* DNA methyltransferases catalyze DNA methylation of neuronal genes during neuronal differentiation and that their deletion results in loss of DNA methylation at multiple neuronal enhancers and aberrant neuronal maturation *in vitro*. Importantly, the results from our *in vitro* analysis further revealed a high efficiency of tamoxifen-induced homozygous deletion of *Dnmt3a* and *Dnmt3b* in NSPCs, making this mouse model suitable for *in vivo* studies of adult hippocampal neurogenesis.

### Deletion of *de novo* DNA methyltransferases in neural precursor cells impairs maturation of new-born neurons *in vivo*

To analyze whether *de novo* DNA methylation is functionally involved in adult hippocampal neurogenesis *in vivo*, we deleted *Dnmt3a* and *Dnmt3b* in NSPCs by tamoxifen administration to nestinCre-ER^T2^::Dnmt3a^(fl/fl)^-Dnmt3b^(fl/fl)^ mice and investigated the functional consequences on adult hippocampal neurogenesis.

We did not observe a difference between WT and KO mice in neural precursor cell proliferation acutely 1 week after tamoxifen administration (Fig. 4A), or in the total number of new-born neurons 5 weeks after induction of deletion (Fig. 4B). To investigate a potential influence on the dynamics of adult neurogenesis, we traced neural precursor cells and their progeny by injecting a lentivirus carrying a Cre-dependent Gfp vector into WT and KO mice (Fig. 4C). After validating the tamoxifen-induced deletion of *Dnmt3a* and *Dnmt3b* in FAC-sorted Gfp-positive cells from KO mice (Fig. S5A), we analyzed cell stage distributions of Gfp-positive cells after a chase period of four weeks. We did not detect a difference in the percentages of label-retaining NSPCs or astrocytes (Fig. S6B-C). Additionally, no differences in total numbers of neural precursor cells and their neuronal progeny were found between KO and WT mice three months after tamoxifen administration (Fig. S6D-G). These results suggested that deleting *de novo* DNA methyltransferases did not result in detectable changes in the neurogenic activity and long-term maintenance of neural precursor cells in the adult hippocampus.

**Figure 4:**
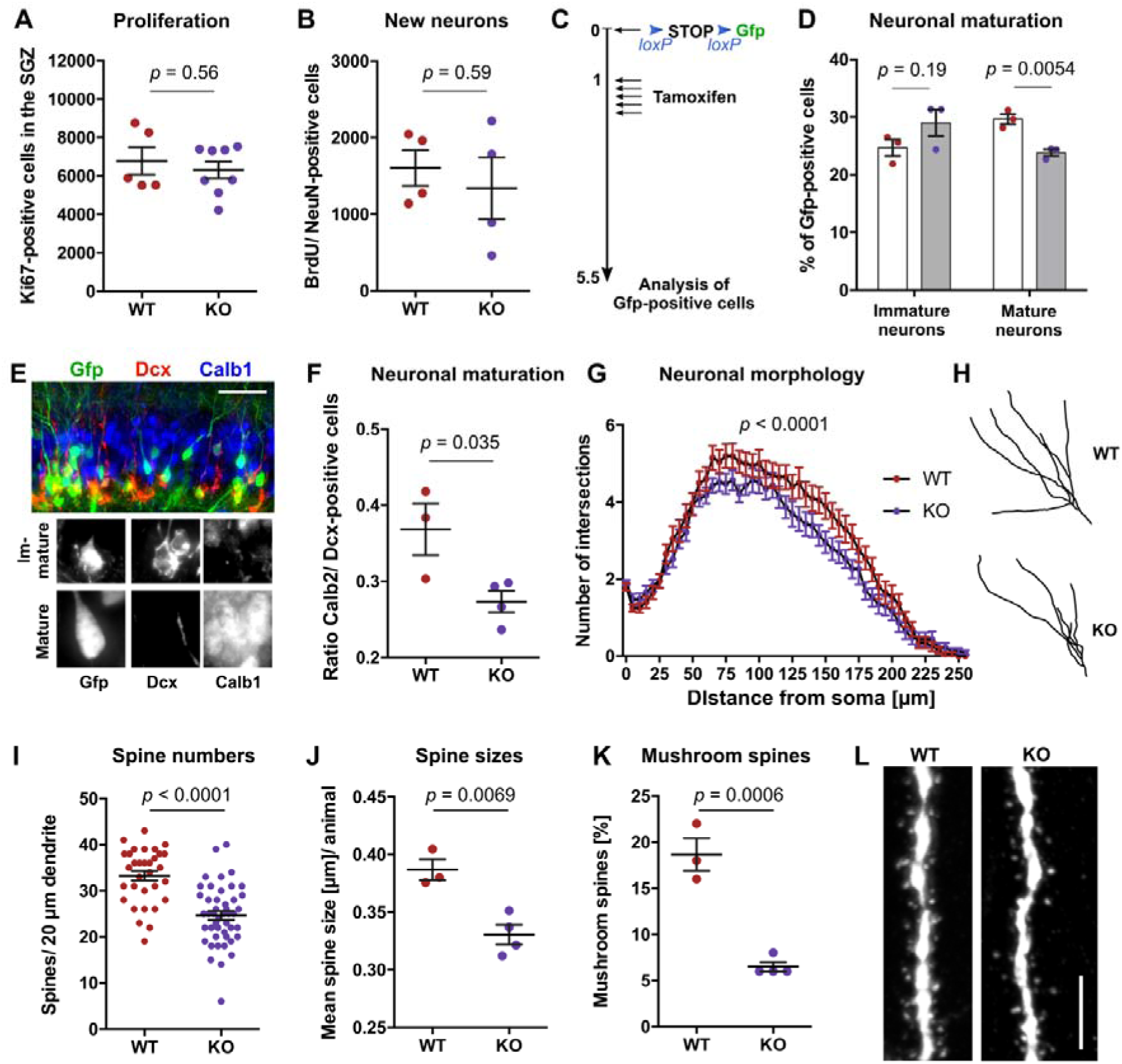
*De novo* DNA methylation controls maturation and synaptogenesis of adult-born neurons in the hippocampus. **A,** No difference in numbers of Ki67-positive, proliferating cells was observed between the subgranular zone (SGZ) of *Dnmt3a*/ *Dnmt3b* WT and *Dnmt3a*/ *Dnmt3b* KO mice one week after tamoxifen administration. **B,** Deletion of *Dnmt3a* and *Dnmt3b* did not change total numbers of new-born neurons. Mice were administered with tamoxifen and analyzed 5 weeks after the first injection. BrdU was injected 3.5 weeks before analysis. **C,** Experimental scheme for results depicted in D-L. Recombined WT and KO cells were identified by Gfp expression. **D,** KO mice show reduced percentages of mature neurons (Calb1-positive, Dcx-negative) among Gfp-positive cells compared to WT mice. Immature neurons were identified as Dcx-positive and Calb1-negative. **E,** Representative fluorescence image for detection of mature and immature neurons. Scale bar: 50 µm. **F,** KO mice show reduced numbers of Calb2-positive cells relative to Dcx-positive cells in the dentate gyrus. **G,** Sholl analysis of Gfp-positive cells revealed reduced dendritic outgrowth of new-born neurons in KO mice. *p*-value from two-way ANOVA. **H,** Representative neuronal traces. **I,** Adult-born neurons of KO mice exhibit reduced numbers of spines in the outer molecular layer. **J,** Mean spine size per animal is reduced in KO mice. **K,** Mean percentage of mushroom spines (diameter of spine head ≥ 0.45 µm) per animal. **L,** Representative fluorescent image of synapses in WT and KO mice. Scale bar: 5 µm.

We next investigated the influence of *de novo* DNA methylation on the dynamics of neuronal maturation. After a chase period of four weeks, KO mice exhibited a significantly lower percentage of mature neurons (Calb1-positive/ Dcx-negative) compared to WT mice but a trend towards increased numbers of immature neurons (Dcx-positive/Calb1-negative; Fig. 4D-E). Similarly, when we compared total numbers of cells expressing the post-mitotic neuronal marker Calb2 with numbers of cells expressing immature neuronal marker Dcx, we detected a significant decrease in the ratio of Calb2-positive cells compared to Dcx-positive cells in KO mice (Fig. 4F), suggesting a delay of neuronal maturation in KO mice. Accordingly, neurons in KO mice showed a mild but significant decrease in dendritic outgrowth compared to neurons in WT mice (Fig. 4G-H).

Synaptogenesis starts three weeks after birth of new hippocampal neurons and is a prerequisite for their experience-dependent activation and functional integration (Toni and Schinder, 2016). Neurons of KO mice exhibited significantly fewer synapses in the outer molecular layer (Fig. 4I). Moreover, spines in KO mice had smaller head diameters (Fig. 4J) and comprised of a lower percentage of mushroom spines (Fig. 4K-L). These results suggest that *de novo* DNA methylation is functionally involved in the maturation and synaptogenesis of new-born neurons in the hippocampus.

### Relationship between *de novo* DNA methylation and transcriptional changes during adult hippocampal neurogenesis

Although DNA methylation has traditionally been implicated in the long-term silencing of gene expression, recent studies suggested that enhancer methylation in neural precursor cells is linked to transcriptional activation (Wu et al., 2010). To investigate whether *de novo* DNA methylation affected neurogenesis-associated transcriptomic changes, we labeled recombined neural precursor cells using a lentivirus (as described in Fig. 4C), isolated Gfp-positive cells from the dentate gyrus of WT and KO mice four weeks after tamoxifen administration and performed single-cell RNA sequencing.

We obtained sufficient sequencing reads for 212 WT cells and 284 KO cells. Visualizing the cells using *t*-distributed Stochastic Neighborhood Embedding (t-SNE) revealed four distinct clusters (Fig. 5A) that corresponded to the different cell stages of adult hippocampal neurogenesis (Fig. 5B). Cluster T1 showed expression of NSPC markers but absence of proliferation markers (Fig. 5B; Fig. S6), identifying it as the quiescent neural stem cell population, while cluster T2 contained the proliferating precursor cells. Cells in cluster T3 corresponded to the postmitotic, neuronally committed precursor cells that expressed *Eomes*. Cluster T4 contained the immature neurons with high expression of markers such as *Dcx*, *Neurod1* and *Calb2*. KO cells showed a similar distribution into clusters as WT cells, but exhibited trends toward higher numbers of immature neurons in cluster 4 (Fig. 5C). Such accumulation of immature neurons in KO mice is consistent with the increased percentage of immature neurons observed in the histological analysis and supports the observed delay in neuronal maturation in KO mice.

**Figure 5:**
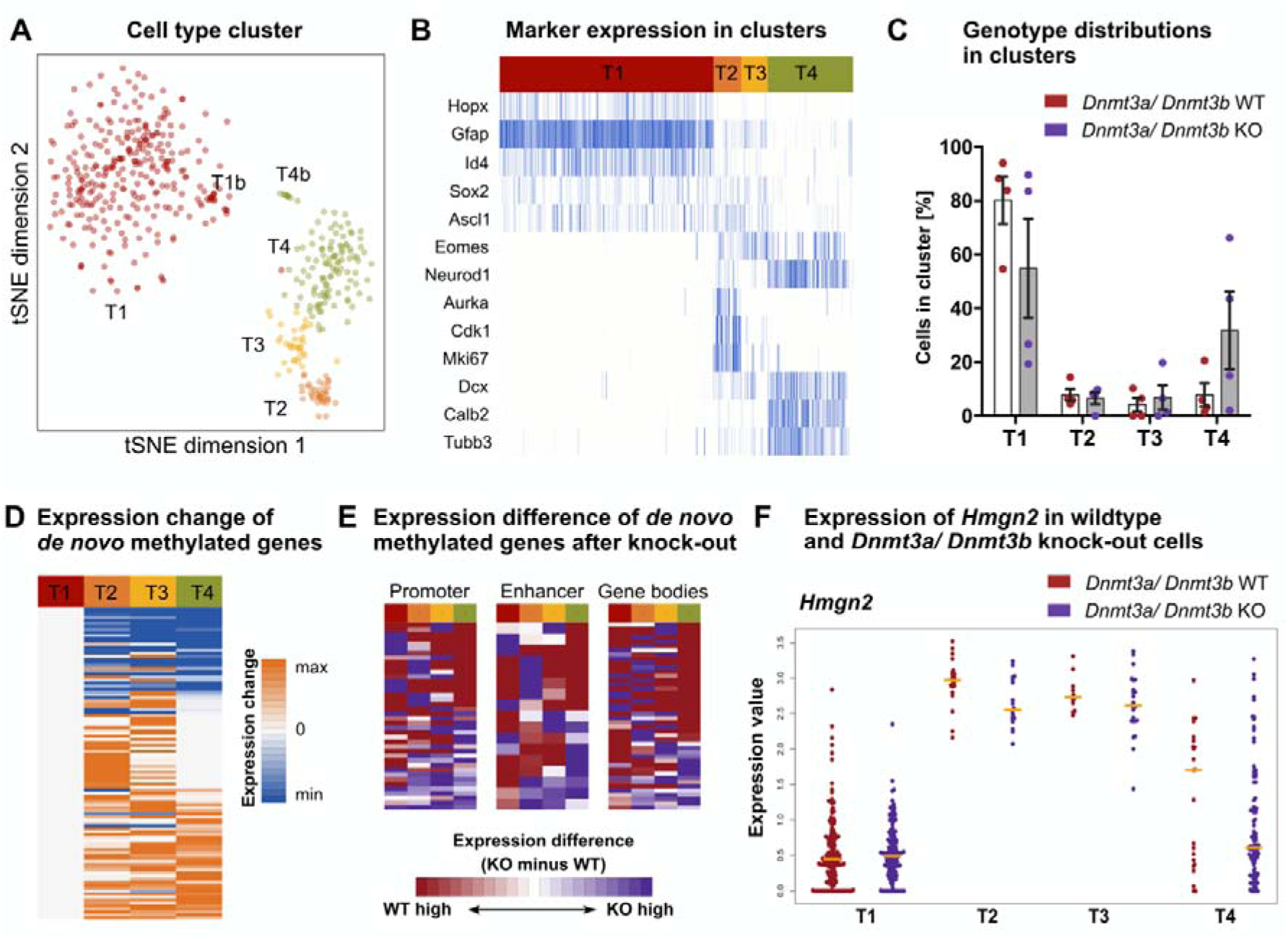
Relationship between neurogenesis-associated *de novo* DNA methylation and transcriptional changes. Experimental outline as depicted in Fig. 4C. Gfp-positive cells were isolated by FACS from WT and *Dnmt3a*/ *Dnmt3b* KO mice and their transcriptional profile analyzed using single-cell RNA sequencing (n = 4 mice per group). **A,** *t*-distributed Stochastic Neighborhood Embedding (t-SNE) clustered cells into 4 populations which are labeled with T1-T4 and highlighted by color. Clusters T1b and T4b were identified as sub-clusters of T1 and T4, respectively. **B,** Marker expression identified the cell clusters as distinct cell stages during adult hippocampal neurogenesis. **C,** Distribution of WT (red) and KO cells (violet) into the different cell clusters. **D,** Predominant increase in expression of neurogenesis-associated *de novo* methylated genes during the neurogenic trajectory in WT cells. Depicted is the relative expression of genes in clusters T2-4 compared to cluster T1. Non-expressed genes were not plotted. **E,** Expression difference of *de novo* methylated genes in KO cells versus WT cells in the individual cell clusters. Genes were separated by genomic location of the DNA methylation change. **F,** Candidate gene showing expression changes between WT and KO cells in the different cell clusters. Individual dots represent expression in a single cell. Orange lines indicate medians.

Despite a chase period of four weeks before analysis, we could not identify mature neurons based on expression of mature neuronal markers, such as Calb1 (Fig. S6), and observed an overrepresentation of precursor cell stages compared to the results from our histological analysis. This can likely be attributed to the rupture of neuronal axons and membranes during the FACS process. We therefore focused our analysis on the DNA methylation-dependent transcriptional changes during early neuronal differentiation in the hippocampus.

To relate neurogenesis-associated DNA methylation changes to transcription, we first analyzed whether *de novo* methylated genes change expression during the differentiation from NSPCs to immature neurons (Fig. 5D). We observed that some of *de novo* methylated genes were down-regulated during the neurogenic trajectory, which would be consistent with a role of DNA methylation in the silencing of stem cell-specific genes during differentiation. However, we unexpectedly found that a substantial part of *de novo* methylated genes increased expression during adult neurogenesis. Among those were genes that showed a transient up-regulation in clusters T2 or T3 as well as genes that were up-regulated only in immature neurons or throughout differentiation. To investigate whether *de novo* DNA methylation drives neurogenesis-associated transcriptional changes, we performed unbiased differential expression analyses between WT and KO cells in the different cell clusters. Although we identified 444 genes that changed expression between the neurogenic cell stages (Supplemental data 3), we could not detect any significantly differentially expressed genes between WT and KO cells. When we plotted the expression difference KO versus WT of *de novo* methylated genes, we found a cluster of genes with higher expression in immature neurons of WT cells (Fig. 5E). An example for such a gene is *Hmgn2* – a transcriptional activator that has been implicated in the control of neurogenesis (Fig. 5F). Together, these results suggest that *de novo* DNA methylation mostly correlated with transcriptional up-regulation during adult neurogenesis, but blocking *de novo* DNA methylation did not cause major transcriptional changes during early neuronal differentiation.

### Neurogenesis-associated *de novo* DNA methylation is required for the experience-dependent activation of the hippocampus

Environmental enrichment (ENR) is a strong pro-neurogenic stimulus that predominantly increases the survival of new-born neurons (Kempermann et al., 1997; Körholz et al., 2018) by mechanisms involving neuronal activation in the hippocampus (Kirschen et al., 2017). We have previously shown that ENR changes DNA methylation patterns at neuronal plasticity genes in the dentate gyrus (Zocher et al., 2020). We here analyzed whether dynamic DNA methylation in NSPCs and immature neurons is required for the pro-neurogenic effect of ENR and for ENR-stimulated hippocampal excitability.

After confirming that housing in ENR for five weeks increases numbers of immature neurons in the dentate gyrus (Fig. S7A), we performed RRBS of FAC-sorted NSPCs, lPCs and neurons from the dentate gyrus after five weeks of ENR and compared them to mice housed in standard housing cages. We detected ENR-induced DNA methylation changes in all three cell populations (Fig. S7B), suggesting that the methylome of different cell populations during adult neurogenesis is susceptible to environmental stimulation. To analyze whether ENR-induced *de novo* DNA methylation is functionally involved in new-born neuron survival, we administered tamoxifen to WT and KO mice and housed them in ENR or STD for five weeks or three months. Deletion of *de novo* DNA methyltransferases did not abolish the ENR-induced increase in the numbers of new-born cells in the hippocampus (Fig. 6A-C), suggesting that *de novo* DNA methylation during adult hippocampal neurogenesis is not required for the pro-neurogenic effect of ENR.

**Figure 6:**
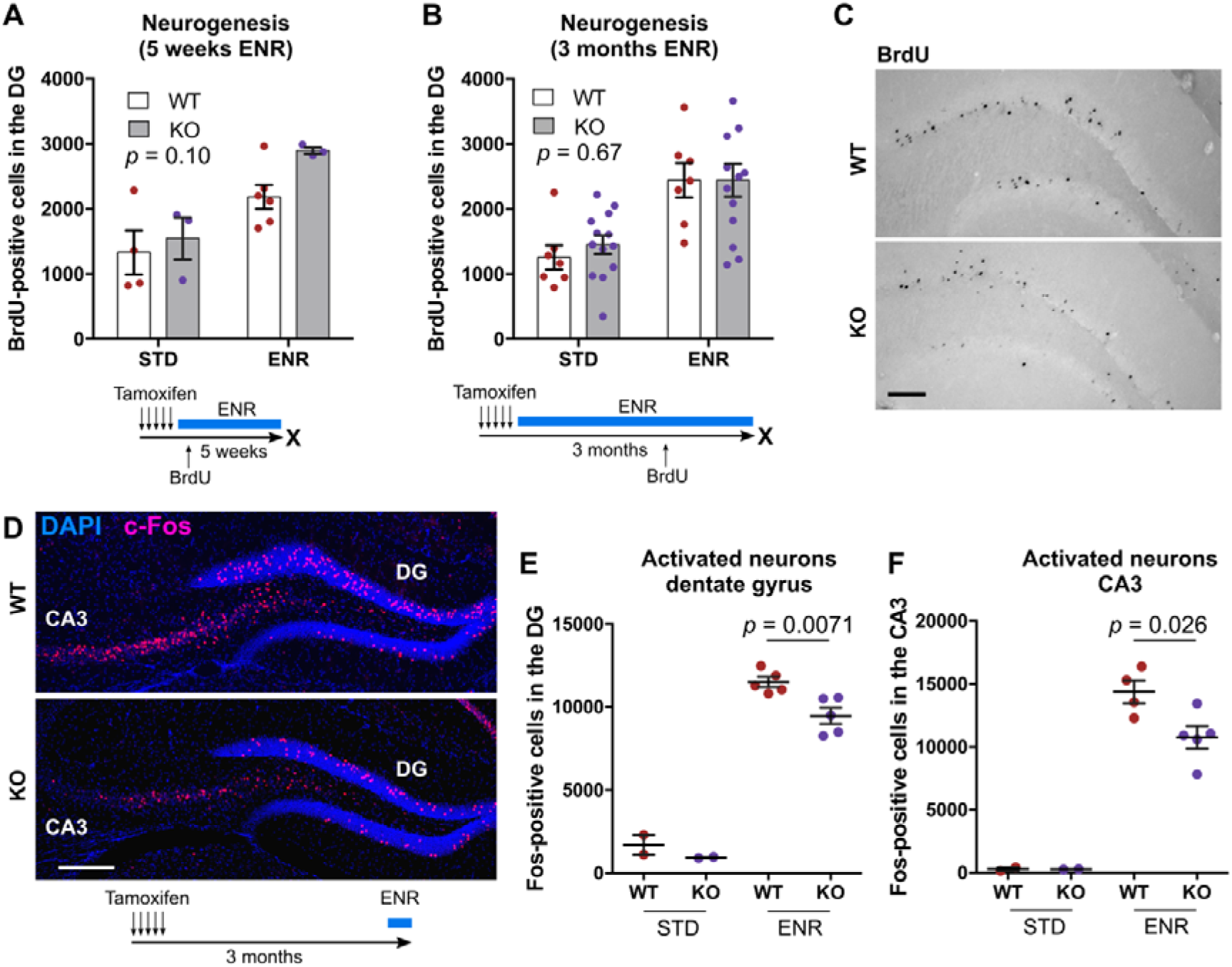
Deletion of *de novo* DNA methyltransferases does not abolish the pro-neurogenic effect of environmental enrichment but impairs hippocampal excitability. A-B, Quantification of adult-born (BrdU-positive) cells in the dentage gyrus (DG) of WT and KO mice after five weeks or three months of environmental enrichment (ENR) or standard housing (STD). BrdU was injected 4.5 weeks before mice were killed. Depicted *p*–values correspond to genotype effects from two-way ANOVA. Influences of ENR were significant with *p* < 0.01 in A-B. **C,** Representative immunohistochemical staining for detection of BrdU-positive cells. Shown are images from WT and KO mice housed in ENR for three months. Scale bar: 100 µm. **D,** Representative fluorescent image of distribution of c-Fos-positive cells in the DG and CA3. Scale bar: 200 µm. **E-F,** Numbers of c-Fos-positive cells in the DG and CA3 without (STD) and with acute stimulation in ENR for 1 h. Depicted *p-*values in E-F are from unpaired *t*-test.

Proper integration of adult-born neurons into the hippocampal circuitry controls neuronal activity in the hippocampus (Berdugo-Vega et al., 2020; Ikrar et al., 2013). To get an idea whether Dnmt3a/b-deficient neurons are impaired in their functional integration, we acutely exposed mice to ENR for one hour and quantified the numbers of activated, c-Fos-positive neurons. KO mice showed significantly fewer c-Fos-positive cells in the dentate gyrus and CA3 of the hippocampus after stimulation in ENR (Fig. 6D-F). These results indicated a critical role for *de novo* DNA methyltransferases for the experience-dependent activation of the hippocampus.

### Neurogenesis-associated *de novo* DNA methylation is crucial for hippocampus-dependent learning and memory

Adult-born neurons of the hippocampus contribute to cognitive flexibility which can be assessed using behavioral tests measuring learning and memory performance. To analyze whether the observed changes in neuronal maturation and hippocampal activation lead to cognitive deficits in KO mice, we tested animals in different behavioral tests. WT and KO mice showed no differences in motor coordination on the rotarod (Fig. S8A-B), exploration of an open field arena (Fig. S8C-D) or novel object recognition (Fig. S8E-F), but a specific impairment in reversal learning in the Morris watermaze (Fig. 7). Previous studies have demonstrated that adult-born hippocampal neurons contribute to the cognitive flexibility required for re-learning of the platform position after reversal of its initial location (Garthe et al., 2009).

**Figure 7:**
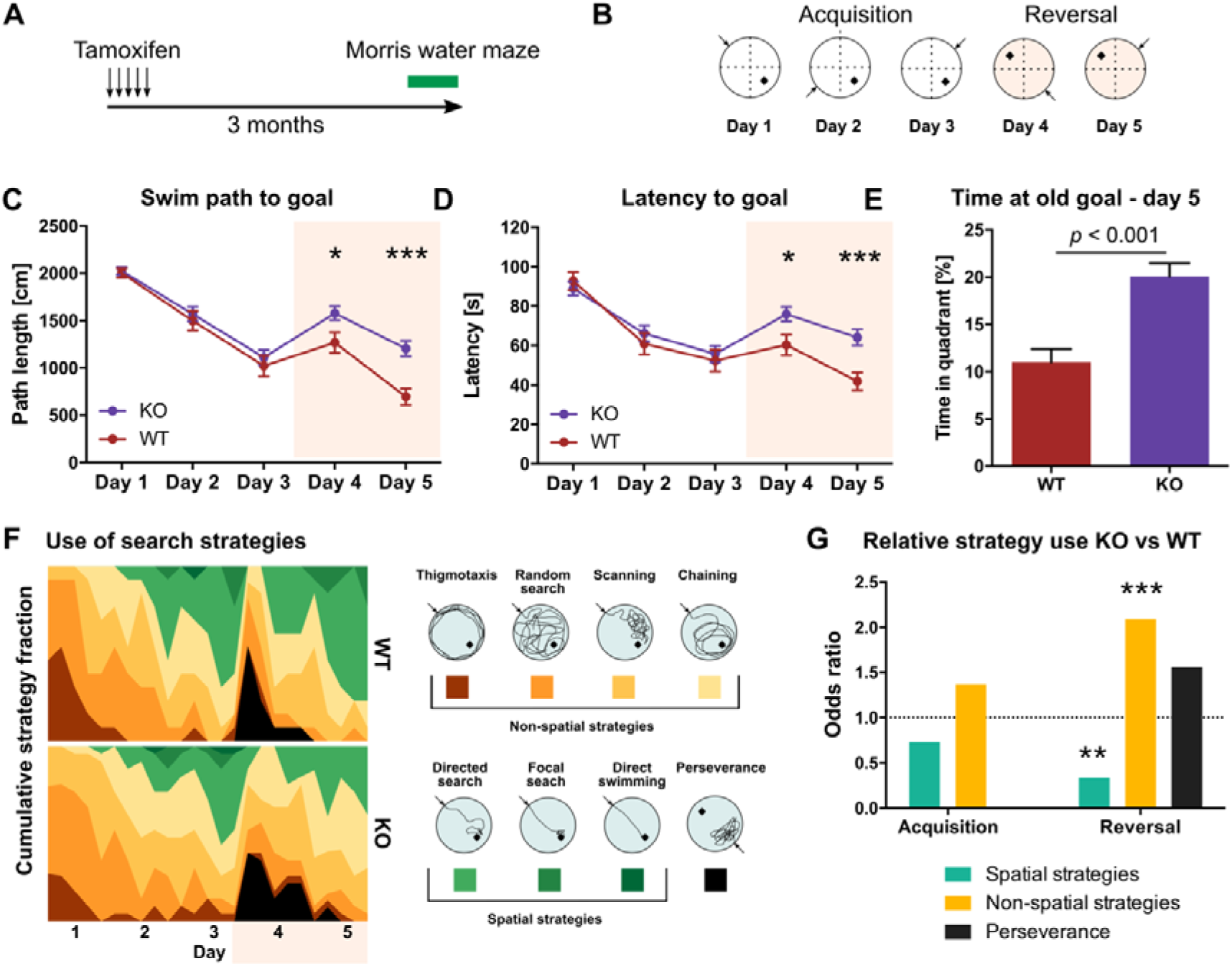
Blocking *de novo* DNA methylation during adult hippocampal neurogenesis impairs hippocampus-dependent learning and memory. **A,** Mice with deletion of *Dnmt3a* and *Dnmt3b* (KO) and controls (WT) were tested in the Morris watermaze three months after administration of tamoxifen. **B,** Protocol for the Morris water maze test. Platform positions are indicated by a black square. Starting positions are highlighted with an arrow. **C,** Average distance animals swam per day to find the hidden platform. **D,** Average time animals need to reach the platform per day. Note that KO mice showed increased path length and latency on days 4 and 5. **E,** Time mice spent on day 5 in the quadrant in which the goal was located during the acquisition phase. **F,** Heatmaps depicting the percentage of individual search strategies used by WT and KO mice per trial (left). Color scheme for the classification into spatial and non-spatial search strategies. **G,** Odds ratio of strategy use in KO compared to WT mice separated by acquisition and reversal phase (*p*-values were determined by fitting a general linear model with binomial distribution). * *p* < 0.05; *** *p* < 0.01; *** *p* < 0.001. (n = 13, WT; n = 23, KO). C-E show mean ± standard error of the mean. Unpaired *t*-test was used to determine significance in C-E.

No difference was detected between WT and KO mice in their ability to learn the initial platform position during the acquisition phase (Fig. 7A-D). However, after reversal of the platform position, KO mice swam significantly longer distances (swim path; Fig. 7C) and needed more time to find the hidden platform compared to WT mice (Fig. 7D). Additionally, on the second day after reversal, KO mice spent more time in the quadrant in which the platform was located before reversal (Fig. 7E). These results suggested that KO mice are less flexible in re-learning the new platform position after reversal. As an additional adult neurogenesis-dependent measure of learning, we analyzed the navigational strategies mice used to allocate the hidden platform. While there was no difference in search strategy choice between KO and WT mice during the acquisition phase, KO mice used significantly fewer hippocampus-dependent “spatial” strategies and more “non-spatial” strategies after reversal (Fig. 7E-F). Together, our results indicate that *de novo* DNA methylation during adult neurogenesis is required for proper functional integration of adult-born neurons into the hippocampus.

## Discussion

Here, we provide a comprehensive characterization of the DNA methylation changes during adult hippocampal neurogenesis and a careful dissection of their functional consequences for adult neural stem cell potential, neuron formation and hippocampal function. We found that, during adult neurogenesis, *de novo* DNA methyltransferases target enhancers and gene bodies of neuronal genes to establish neuron-specific methylomes and expression patterns. Our single-nucleotide resolution DNA methylation data of isolated NSPCs, lPSCs and neurons provides the first genomic targets of epigenetic changes during *in vivo* adult neurogenesis and will be a unique resource for the field. Our mechanistic analyses demonstrated a critical role of neurogenesis-associated *de novo* DNA methylation specifically for the maturation of new-born neurons, hippocampal excitability and for the cognitive enhancement conveyed by adult hippocampal neurogenesis. Together, our study provides new insight into the role of *de novo* DNA methylation for an adult stem cell system that influences life-long behaviour and cognition.

The low numbers of NSPCs in the adult hippocampus has so far hampered genome-wide epigenetic analyses. Using an optimized protocol for reduced representation bisulfite sequencing, we here show the pronounced *de novo* DNA methylation of neuronal genes during differentiation of adult NSPCs into hippocampal neurons. A predominant gain in DNA methylation has also been found during the *in vitro* neuronal differentiation from cortical progenitor cells of the embryonic brain (Luo et al., 2019), during differentiation of embryonic stem cells into motor neurons (Ziller et al., 2018) or pyramidal neurons (Mohn et al., 2008) and during the reprogramming of fibroblasts into neurons (Luo et al., 2019). How much of the detected neuronal *de novo* DNA methylation is attributed to 5-hydroxymethylcytosine rather than 5-methylcytosine is currently unknown, but its future investigation could help to further resolve the link to gene expression. Previous studies of embryonic neurogenesis reported a specific increase in 5-hydroxymethylcytosine at neurogenic genes which correlated with transcriptional up-regulation (Hahn et al., 2013; Noack et al., 2019). Our single-cell RNA sequencing data suggested that increases in DNA methylation during adult neurogenesis occurred predominantly in proximity to genes which increased gene expression. This is in accordance with the detected enrichment of *de novo* DNA methylation at neuronal and synapse-related genes and consistent with findings of Wu and colleagues who showed that Dnmt3a-dependent DNA methylation of neuronal genes activated transcription through repression of polycomb proteins (Wu et al., 2010). Moreover, many pro-neuronal transcription factors specifically bind to methylated genomic sites, such as Klf4 – which was enriched at *de novo* methylated enhancers – but also Pax6, Nfatc4, Npas2, Nr2f1, Tfcp2, Egr4 and Snai1 (Yin et al., 2017). Our DNA methylation data will be an important starting point for future experiments to understand how *de novo* DNA methyltransferases are recruited to neuronal genes and how they interact with other molecular players to control adult hippocampal neurogenesis.

Our study suggests that neurogenic commitment is epigenetically pre-determined in adult neural stem cells and independent of further *de novo* DNA methylation during neuronal differentiation. Previous studies showed that differentiation of embryonic stem cells into NSPCs is associated with pronounced *de novo* DNA methylation (Mohn et al., 2008; Ziller et al., 2018). Blocking these changes through deletion of *Dnmt3a* and *Dnmt3b* in embryonic stem cells significantly impaired the terminal differentiation of neural progenitor cells into neurons, resulting in increased neuronal cell death and reduced percentages of neuronal progeny but enhanced astrogliogenesis (Wu et al., 2012; Ziller et al., 2018). Similarly, mice with global knock-outs of *Dnmt3a* showed reduced postnatal neurogenesis in the hippocampus and subventricular zone (Wu et al., 2010). In our study, deleting *Dnmt3a* and *Dnmt3b* specifically in adult NSPCs did not impair the *in vivo* production of new neurons at the level of NSPCs proliferation, neuronal fate specification or survival. This suggests that *de novo* DNA methylation associated with the terminal differentiation of NSPCs into neurons are not required for making a neuron *per se*. Neuronal fate determination might be dependent rather on the maintenance of the previously established DNA methylation patterns in NSPCs than on continuous activity of *de novo* DNA methyltransferases during terminal differentiation. Consistent with this model, inducible deletion of *Dnmt1* in adult NSPCs, which results in a proliferation-dependent loss of DNA methylation in stem cells (Liao et al., 2015), strongly reduced numbers of new-born neurons in the hippocampus, while it left proliferation or long-term maintenance of NSPCs unaffected (Noguchi et al., 2015).

Rather than controlling mere numbers of new-born neurons in the hippocampus, *de novo* DNA methyltransferases were crucial for the morphological and functional maturation of the new neurons. Evidence of whether this deficit is a direct consequence of hampered *de novo* DNA methylation during neurogenesis or whether mature neurons need continuous Dnmt3a/b-dependent epigenetic remodeling would require additional cell stage-specific deletion of Dnmt3a/b during adult neurogenesis. However, the strong enrichment of *de novo* DNA methylated CpGs at synaptic plasticity-related genes are suggestive of a direct role of *de novo* DNA methyltransferases in controlling neuronal maturation. Moreover, the reduced excitability we found in the dentate gyrus and CA3 of KO mice after acute stimulation supports a direct role of *de novo* DNA methyltransferases in the control of functional maturation. New-born neurons have distinct electrophysiological properties compared to mature granule cells (Ge et al., 2007; Gu et al., 2012) and alter the excitability of the hippocampus, through mechanisms such as feed-forward inhibition and others (Berdugo-Vega et al., 2020; Ikrar et al., 2013; Jakubs et al., 2006). Understanding how Dnmt3a/b-deficient adult-born neurons impair hippocampal excitability – whether through altered input sensitivity, quality of their firing pattern, altered neuronal connectivity or a combination of other mechanisms – could help, in future experiments, unraveling the mechanisms of how adult-born neurons promote cognitive flexibility.

ENR changed DNA methylation patterns in neural precursor cells of the adult hippocampus, but deleting *de novo* DNA methyltransferases during neurogenesis did not abolish ENR-induced increases in adult neurogenesis. Although these results show that cell-intrinsic *de novo* DNA methylation are not required for the enhanced survival of new-born neurons, they do not generally exclude a functional role of ENR-induced DNA methylation changes in mediating brain plasticity. First, instead of controlling cell numbers, *de novo* DNA methylation could mediate the previously reported ENR-induced acceleration of new-born neuron maturation (Alvarez et al., 2016; Bergami et al., 2015). Second, ENR led to a predominant hypomethylation of neurons, which is consistent with the previously reported hypomethylation of the dentate gyrus in response to neuronal activation (Guo et al., 2011). Tet enzymes are responsible for active DNA demethylation and have been shown to control numbers of new neurons in the adult hippocampus (Gontier et al., 2018; Zhang et al., 2013). Hence, ENR-induced hypomethylation through Tet enzyme activity could potentially mediate ENR-induced increases in adult hippocampal neurogenesis. Moreover, survival of new neurons in the hippocampus is regulated by local network activity (Song et al., 2016), which is influenced by DNA methyltransferases (Feng et al., 2010). Therefore, ENR-induced DNA methylation changes in mature neurons rather than cell-intrinsic DNA methylation changes in neural precursor cells and new-born neurons could mediate the net increase in adult hippocampal neurogenesis. Investigating the functional contribution of ENR-induced DNA methylation changes for activity-dependent brain plasticity and behavior will be an exciting area for future research toward understanding of how experiences shape brain activity.

Proper control of neuronal DNA methylation patterns is necessary for life-long brain function (Gontier et al., 2018; Oliveira et al., 2012) and its dysregulation is a proposed mechanism underlying several neurological diseases, including Alzheimer’s disease and depression (Han et al., 2018; Li et al., 2019). Here, we show that *de novo* DNA methyltransferases are required for the contribution of adult neurogenesis to hippocampus-dependent learning and memory. Our study provides evidence for a role of *de novo* DNA methylation for adult neural stem cell potential and a resource for further mechanistic studies toward understanding the molecular control of healthy brain function.

## Acknowledgements

This study was financed from basic institutional funds (Helmholtz Association and TU Dresden). SZ was supported by a fellowship from the International Max Planck Research School on the Life Course, Berlin, and an EMBO Short term fellowship. The authors thank the “DRESDEN-concept Genome Center” for excellent sequencing service. We further thank Sandra Günther and Jens Bergmann for animal care as well as Silke White and the DZNE imaging facility for support with microscopy. We are grateful to Tomohisa Toda and Florian Noack for their comments on the manuscript, to Stefan Hans for sharing the 2lox-Gfp plasmid and to Jadna Bogado Lopes for technical support.

## Author contributions

SZ conceptualized the study, performed the experiments, analyzed the data and wrote the manuscript. FC, GBV and GK reviewed and edited the manuscript. RWO analyzed single-cell RNA sequencing data. GBV prepared viruses and performed stereotactic injections. NR and VSA performed FACS. AK coordinated breeding of mice. CG performed genotyping of mice and cell lines. NR and SS helped with immunohistochemistry. KH and JLS performed single-cell RNA sequencing. FC supervised virus injections. GK supervised the study and acquired funding.

## Methods

### Animals

Female C57BL/6JRj mice were purchased from Janvier at an age of five weeks. *Nestin*::Gfp (Yamaguchi et al., 2001), *Dcx*::Gfp mice (Gong et al., 2003) and *nestin*::CreER^T2^- Dnmt3a^(fl/fl)^-Dnmt3b^(fl/fl)^ mice were bred and maintained at the animal facility of the Center for Regenerative Therapies Dresden. To generate *nestin*::CreER^T2^-Dnmt3a^(fl/fl))^-Dnmt3b^(fl/fl)^ mice, Dnmt3a-2lox (Kaneda et al., 2004) and Dnmt3b-2lox (Dodge et al., 2005b; both obtained from RIKEN, Japan) were crossed and the offspring mated with *nestin*::CreER^T2^ mice (Imayoshi et al., 2006). Experimental animals were obtained by backcrossing *nestin*::CreER^T2(+/-)^-Dnmt3a^(fl/fl)^-Dnmt3b^(fl/fl)^ mice with *nestin*::CreER^T2(-/-)^-Dnmt3a^(fl/fl)^- Dnmt3b^(fl/fl)^ mice. WT animals (*nestin*::CreER^T2(-/-)^-Dnmt3a^(fl/fl)^-Dnmt3b^(fl/fl)^) were the siblings of the KO animals that did not contain the *nestin*::CreER^T2^ allele. For lineage tracing experiments, WT animals were *nestin*::CreER^T2(+/-)^-Dnmt3a^(wt/wt)^-Dnmt3b^(wt/wt)^ and derived from the offspring of the original cross of Dnmt3a^(fl/wt)^-Dnmt3b^(fl/wt)^ with *nestin*::CreER^T2^. The parental strains Dnmt3a-2lox, Dnmt3b-2lox and *nestin*::CreER^T2^ were maintained on C57BL/6 backgrounds.

Mice were housed in standard polycarbonate cages (Type II, Tecniplast) and maintained on a 12 h light/dark cycle at the animal facility of the Center for Regenerative Therapies Dresden and the German Center for Neurodegenerative Diseases. Food and water were provided *ad libitum*. All experiments were conducted in accordance with the applicable European and National regulations (Tierschutzgesetz) and were approved by the local authority (Landesdirektion Sachsen: file numbers 25-5131/354/63 and 25-5131/365/9).

### Administration of drugs

Bromodeoxyuridine (BrdU; Sigma) was dissolved in 0.9 % sodium chloride and injected intraperitoneally at a final concentration of 50 mg/kg body weight. Tamoxifen (Sigma) was dissolved in corn oil (Sigma) and 250 mg/kg body weight were administered via oral gavage (one injection every day for five consecutive days).

### Fluorescence-activated cell sorting of dentate gyrus tissue

Micro-dissected dentate gyri were cut into pieces with a scalpel blade and enzymatically dissociated using the Neural Tissue Dissociation Kit P (Miltenyi Biotec) according to the manufacturer’s manual. Dissociated cells were washed with Hank’s balanced salt solution (HBSS; Gibco) and separated using a 40 µm cell strainer (Falcon, BD Bioscience). To remove dead cells, propidium iodide (1 µg/ml; Thermo Fisher Scientific) was added directly before the sort. For labeling of neurons, cells were incubated with NeuroFluor^TM^ NeuO (1:500; Stem Cell Technologies) for 15 min at 4 °C in HBSS. Fluorescence-activated cell sorting (FACS) was performed using a BD FACSARIA^TM^ III sorter and the software FACSDiva v 8.0.1 (both BD Bioscience). Data was analyzed using FlowJo v 10 (Tree Star, Inc.).

### Reduced Representation Bisulfite Sequencing

Genomic DNA was isolated using the QIAamp DNA Micro Kit (QIAGEN). RRBS libraries were prepared using the Premium RRBS Kit (Diagenode) and purified twice using Agencourt AMPure XP beads (Beckman Coulter; 1X bead volume). Each library from NSPCs and lPCs was generated with a start material of 8,000 cells (pools of three mice per sample). Libraries from *in vitro* and *in vivo*-derived neurons were prepared from 100 ng genomic DNA. Quality and concentration of RRBS libraries were determined using the High Sensitivity NGS Fragment Analysis Kit (Advanced Analytical) and a fragment analyzer with capillary size of 33 cm (Advanced Analytical). Sequencing was performed using a HiSeq2500 or a NextSeq500 platform in a 75 bp single end mode with a minimum sequencing depth of 10 million reads per sample.

### Quantitative polymerase chain reaction

Total RNA was isolated using the RNeasy Micro Kit (QIAGEN). RNA was reversely transcribed into complementary DNA using SuperScript^TM^ II Reverse Transcriptase with 500 ng Oligo(dT)_12-18_ primers and 1 μl dNTPs (10 mM each; all Thermo Fisher Scientific). Quantitative real-time polymerase chain reactions were performed using the QuantiFast SYBR Green PCR Kit (QIAGEN) and the CFX ConnectTM Real-Time PCR Detection System (Bio-Rad). Primers for quantitative PCRs are listed in Supplementary Table 1.

**Supplementary Table 1.**
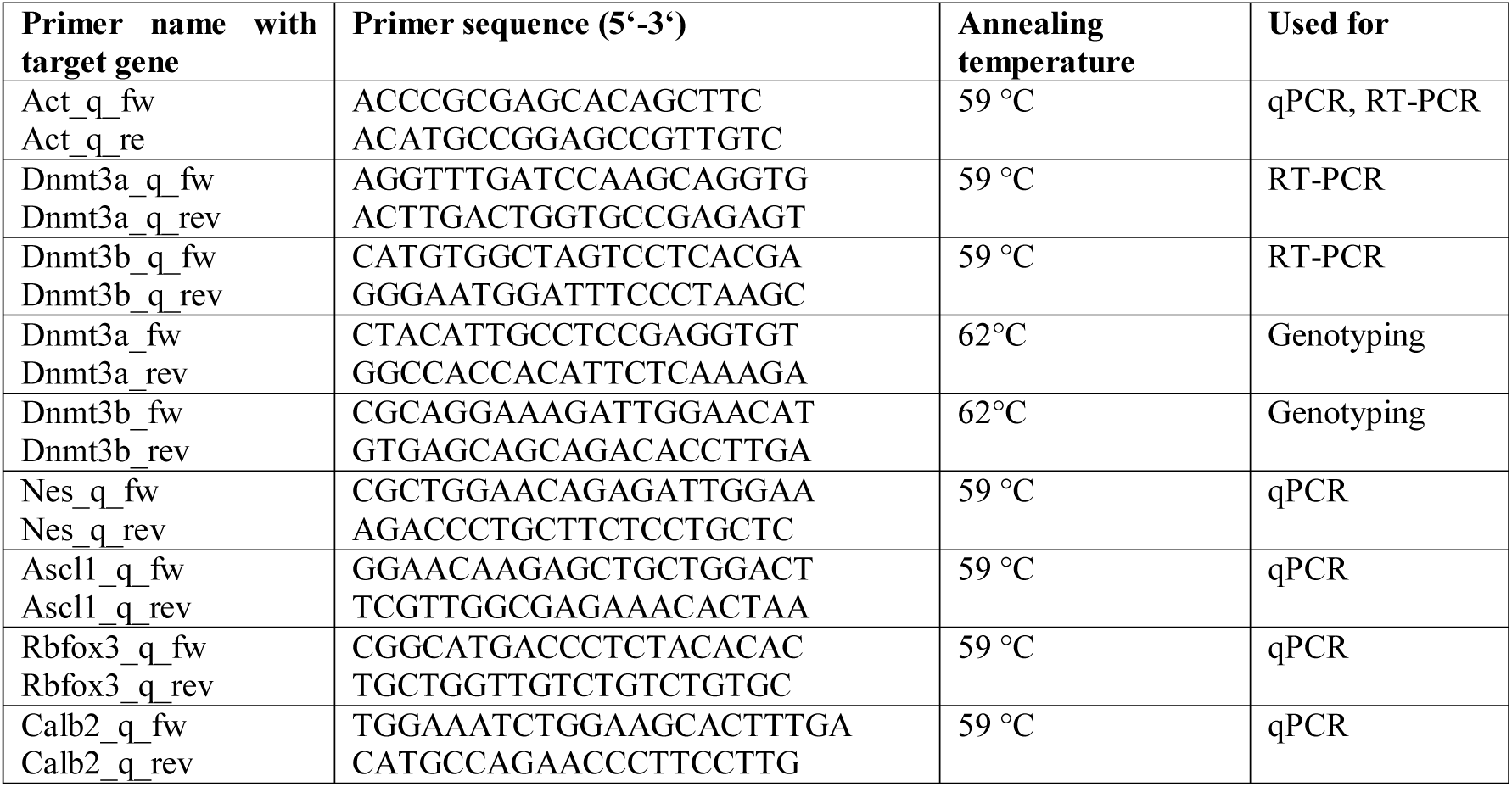
List of primer sequences.

### Single-cell RNA sequencing

Single cells were collected into a 384-well plates pre-loaded with 2.3 µl lysis buffer (50 mM guanidine hydrochloride, 17.4 mM dNTPs, 2.2 µM SMART dT30VN primer). Cells from WT and KO mice were randomized across 3 individual plates. The plates were sealed and stored at −80 °C. Smart-Seq2 libraries were generated using Tecan Freedom EVO and Nanodrop II (Bionex) systems as previously described (Picelli et al., 2013).

In brief, cells were incubated at 95 °C for 3 min. Reverse transcription was performed by adding 2.7 µl reverse transcription mix (SuperScript II buffer (Invitrogen), 9.3 mM DTT, 370 mM Betaine, 15 mM MgCl_2_, 9.3 U SuperScript II RT (Invitrogen), 1.85 U recombinant RNase Inhibitor (Takara), 1.85 µM template-switching oligo) to each lysed cell using a Nanodrop II liquid handling system (BioNex) followed by incubation at 42 °C for 90 min and inactivation at 70 °C for 15 min. Amplification of cDNA was performed for 16 cycles with 7.5 µl preamplification mix containing KAPA HiFi HotStart ReadyMix and 2 µM ISPCR primers. cDNA was purified with 1X Agencourt AMPure XP beads (Beckman Coulter) and eluted in 14 µl nuclease-free water. Concentration and cDNA size was analyzed for selected representative wells using a High Sensitivity D5000 assay for the Tapestation 4200 system (Agilent). cDNA was diluted to an average of 200 pg/µl and 100 pg cDNA from each cell and was tagmented using Nextera XT DNA Library Preparation Kit (Illumina). The tagmentation reaction was incubated at 55 °C for 8 min before removing Tn5 from the DNA by adding 0.5 µl NT buffer per well. 1 µl well-specific indexing primer mix from Nextera XT Index Kit v2 Sets A-D and 1.5 µl NPM was added to each well and the tagmented cDNA was amplified for 14 cycles according to manufacturer’s specifications. PCR products from all wells were pooled and purified with 1X Agencourt AMPure XP beads (Beckman Coulter). The fragment size distribution was determined using a High Sensitivity D5000 assay for the Tapestation 4200 system (Agilent) and library concentration was determined using a Qubit dsDNA HS assay (Thermo Fischer). Libraries were clustered at 1.4 pM concentration using High Output v2 chemistry and sequenced on a NextSeq500 system SR 75bp with 2*8bp index reads.

### Preparation and administration of lentiviruses

Lentiviruses were produced by polyethyleneimine co-transfection of 293T cells with the transfer vector encoding for GFP, HIV-1 gag/pol and VSV-G as previously described (Artegiani et al., 2012; Berdugo-Vega et al., 2020). The transfer vector was designed such that removal of the floxed-stop codon after tamoxifen-dependent recombination allowed for the expression of the GFP reporter specifically in the NSC progeny. One day after transfection, cells were switched to serum free medium and 1 day later the supernatant was filtered and centrifuged at 25,500 rpm for 4 h. The viral particles were resuspended in 40 μl of PBS per 10 cm petri dish and further concentrated using centrifugal filters (Amicon) yielding ca. 40 μl of virus suspension per construct with a titer of 10^8^-10^9^ IU/ml as assessed on HEK cells. Viral particles (1 μl) were stereotaxically injected in the DG hilus of isofluorane-anaesthetized mice as previously reported (Artegiani et al., 2012) using a nanoliter-2000 injector (World Precision Instruments) and a stereotaxic frame Model 900 (Kopf Instruments) at ±1.6 mm mediolateral, –1.9 anteriorposterior, and –1.9 mm dorsoventral from bregma with a constant flow of 200 nl/min.

### Immunohistochemistry

Mice were anesthetized by intraperitoneal injection of a mixture of ketamine (100 mg/kg bodyweight; WDT) and xylazin (10 mg/kg bodyweight; Serumwerk Bernburg AG) diluted in 0.9 % sodium chloride. Mice were transcardially perfused with 0.9 % sodium chloride. Brains were removed from the skull and post-fixed in 4 % paraformaldehyde prepared in phosphate buffer (pH 7.4) overnight at 4 °C. For cryoprotection, brains were transferred into 30 % sucrose in phosphate buffer until they sank down before they were cut into 40 µm- or 60 µm- thick coronal sections using a dry-ice-cooled copper block on a sliding microtome (Leica, SM2000R). Sections were stored at −20 °C in cryoprotectant solution (25 % ethyleneglycol, 25 % glycerol in 0.1 M phosphate buffer, pH 7.4).

For immunofluorescent stainings, free-floating sections were washed in phosphate buffered saline (PBS) and unspecific antibody binding sites were blocked by incubating sections in PBS supplemented with 10 % donkey serum (Jackson ImmunoResearch) and 0.2 % Triton X-100 (Carl Roth) for 90 min at room temperature. Primary antibodies were applied overnight at 4 °C and secondary antibodies for 2 h at room temperature. Antibodies were diluted as depicted in Supplementary Table 2 in PBS supplemented with 3 % donkey serum and 0.2 % Triton X-100. Nuclei were labeled with Hoechst 33342 (1:4000; Jackson ImmunoResearch). Sections were mounted onto glass slides and cover-slipped using Aquamount (Polysciences Inc.). Fluorescence stainings were imaged using a Zeiss Apotome equipped with an AxioCam MRm camera and the software AxioVision 4.8 (Zeiss).

**Supplementary Table 2.** List of antibodies.

| Antibody target | Antibody description | Manufacturer | Catalog number | Dilution |
| --- | --- | --- | --- | --- |
| BrdU | Monoclonal rat | AbD Serotec | - | 1:500 |
| Calbindin (Calb1) | Monoclonal mouse | Swant | 300 | 1:500 |
| Calretinin (Calb2) | Polyclonal goat | Swant | CG1 | 1:500 |
| c-Fos | Polyclonal rabbit | Synaptic Systems | 226003 | 1:1000 |
| Dcx | Polyclonal goat | Santa Cruz | sc-8066 | 1:500 |
| Gfap | Polyclonal rabbit | DakoCytomation | Z0334 | 1:1000 |
| Gfp | Chicken | Abcam | ab13970 | 1:750 |
| Ki67 | Polyclonal rat | eBioscience | 14-5698-82 | 1:750 |
| S100b | Polyclonal rabbit | Abcam | ab52642 | 1:500 |
| Sox2 | Polyclonal goat | R&D | AF2018 | 1:500 |
| Tubb3a | Monoclonal mouse | Promega | G712A | 1:1000 |

BrdU-labeled cells were stained using peroxidase method as previously described (Körholz et al., 2018). Briefly, sections were incubated in 0.6 % hydrogen peroxide for 30 min to inhibit endogenous peroxidase activity. For antigen retrieval, sections were incubated in pre-warmed 2.5 M hydrochloric acid for 30 min at 37 °C. Blocking and antibody incubation was performed as described for immunofluorescent stainings. For detection, Vectastain ABC-Elite reagent (9 µg/ml of each component; Vector Laboratories LINARIS) was used with diaminobenzidine (0.075 mg/ml; Sigma) and 0.04 % nickel chloride. All washing steps were performed in tris buffered saline. Mounted sections were cleared with Neo-Clear and cover-slipped using Neo-Mount (both Millipore). BrdU-positive cells were counted using a brightfield microscope (Leica DM 750).

### Analysis of dendrites and synapses

Imaging was performed using a Confocal LSM 780 NLO and the software ZEN 2011 SP7. Dendritic lengths in the hippocampus were analyzed on 60 µm-thick sections with a 20x objective and 1.7 µm stack intervals. *In vitro* differentiated neurons were imaged using an Apotome and a 20x objective. Sholl analysis was performed in Fiji using the plugin Simple Neurite Tracer with a radius step size of 5 µm. Synapses were quantified on images taken from 40 µm thick sections with a 63x glycerol objective, using 0.8 µm stack intervals and 1.8 x zoom.

### Culture and analysis of *in vitro* neural precursor cells

Micro-dissected dentate gyri were enzymatically dissociated using the Neural Tissue Dissociation Kit P (Miltenyi Biotec). Cells were separated using a 40 µm strainer, plated at approximate single-cell dilution in 96-well plates in neural precursor cell medium supplemented with 20 mg/ml Heparin (MP Biomedicals) and incubated for 14 days. Neural precursor cell medium consisted of Neurobasal medium (Gibco^TM^) supplemented with 1x B-27 supplement (Gibco^TM^), 1x GlutaMAX^TM^ supplement (Gibco^TM^), 10,000 U/ml Pen/Strep (Gibco^TM^), 20 mg/ml human FGF2 (PeproTech) and 20 mg/ml human EGF (PeproTech). For generation of clonal lines, single neurospheres were transferred into 96-well plates, mechanically disrupted using a 200 µm pipet tip and expanded as adherent monolayer cultures. Cells were maintained on dishes coated with poly-D-lysine (Sigma-Aldrich) and laminin (Roche Diagnostics). Neural precursor cells were passaged at a confluence of 80 % using Accutase (Gibco^TM^). Differentiation was induced by withdrawing human EGF and FGF from the neural precursor cell medium for five days.

For genotyping of clonal cell lines, genomic DNA was isolated using the QIAamp DNA Micro Kit (QIAGEN) and gene fragments were amplified using DreamTaq DNA Polymerases (Thermo Fisher Scientific). For calculation of knock-out efficiency, a minimum of ten cell lines from each of the five animals were expanded and genotyped. Five cell lines with homozygous knock-outs of *Dnmt3a* and *Dnmt3b* were randomly selected for characterization.

For proliferation assays, cells were incubated with 10 µM EdU for 3 h and analyzed using Click-iT^TM^ EdU Alexa Fluor 647 Flow Cytometry Assay Kit (Thermo Fisher Scientific). Cells were recorded with a BD^TM^ LSRII (BD Biosciences) and analyzed using FlowJo v10^TM^.

For analysis of cell differentiation, cells were cultured on poly-D-lysine/ laminin coated glass coverslips and fixed with 4 % paraformaldehyde for 10 min. Cells were washed in PBS and blocked for 90 min with PBS supplemented with 10 % normal donkey serum (Jackson ImmunoResearch) and 0.2 % Triton X-100 (Carl Roth). Primary antibodies were incubated for 1 h and secondary antibodies (1:1000; Jackson ImmunoResearch) for 30 min in PBS supplemented with 3 % normal donkey serum and 0.2 % Triton X-100. Nuclei were stained with Hoechst 33342 (1:4000 in PBS) for 10 min. All incubation steps were performed at room temperature.

### Animal behavior

#### Environmental enrichment

The enriched environment was designed such that it provided sensory, physical, social and cognitive stimulation to the mice. The enclosure was made of white polycarbonate walls and covered an area of 0.74 m^2^. For cognitive stimulation, the cage was equipped with tunnels, plastic toys and hide-outs, which were rearranged twice per week. To increase social complexity, at least ten mice were housed at the same time in one enclosure. Dirty bedding material and toys were replaced once per week. Control mice stayed in standard polycarbonate cages (Type II, Tecniplast).

#### Rotarod

Mice were placed on an Economex Rotarod (Columbus Instruments) and trained on three consecutive days with three trials per day. The rotating cylinder started with a speed of 4 rpm and accelerated by 0.1 rpm. Time until mice fell off was measured.

#### Open field and novel object exploration test

Open field and novel object exploration tests were performed as previously described (Körholz et al., 2018) with few modifications. Briefly, mice were placed in a square arena (50 cm^2^) and their movements were recorded for five minutes using Ethovision XT (Noldus). For the novel object recognition test, two identical objects were placed in the arena and the time mice spent around the objects was analyzed. On the fourth day of testing, one of the objects was replaced with a novel object. Objects and novel object location were randomized among mice.

#### Morris watermaze

Mice were tested in the reference memory version of the Morris watermaze test as previously described (Garthe et al., 2009). Briefly, a pool with a diameter of 2 m was filled with opaque colored water that was equilibrated to room temperature (20-22 °C). A quadratic platform with a surface of 10 cm^2^ was placed in the pool hidden under the water surface. Mice were tested on five consecutive days with five trials per day and an inter-trial interval of 1-2 h. In every trial, mice were allowed to search for the hidden platform for 120 s and were placed on the platform afterwards for 15 s irrespective of trial outcome. Starting positions changed every day, but remained constant for all trials on one day. After the third testing day, the platform position was changed to the opposite quadrant. Swim paths and latencies were recorded using a camera (Logitech) and the software EthoVision XT (Noldus). Search strategies were manually scored from swim tracks with the experimenter blinded to the groups. Differences in strategy use were analyzed by fitting a general linear model with binomial distribution using the R function glm. Odds ratios are the exponential coefficients of the model.

### Bioinformatic data analysis

#### DNA methylation data

Fastq reads were trimmed using Trim Galore 0.4.4 and the function *Cutadapt* 1.8.1 in RRBS mode and mapped against mm10 using Bismark 0.19.0 (Krueger and Andrews, 2011). Detection of differentially methylated cytosines was performed using methylKit v1.12.0 (Akalin et al., 2012). Briefly, methylation levels were extracted from sorted Binary Alignment Map files using the function *processBismarkAln*. Data was filtered for cytosines with a minimum coverage of ten reads and a maximum coverage of 99.9 % percentile in at least three samples per group using the functions *filterByCoverage* and *unite*. Differentially methylated cytosines were identified using the *methDiff* function applying the chi-squared test, basic overdispersion correction and multiple testing correction using the Sliding Linear Model (SLIM) method, a significance threshold of *q* < 0.05 and a threshold for absolute cytosine methylation differences greater than 20 %.

Genomic locations of CpG islands, enhancers, exons and introns were retrieved from the UCSC Genome Browser (Karolchik et al., 2004). Promoters were defined as 500 bp upstream and downstream of transcription start sites. Promoter regions were excluded from gene bodies. Overlaps of cytosines was performed using the function *subsetByOverlaps* of the R package Genomic Ranges v1.38.0 (Lawrence et al., 2013).

Differentially methylated cytosines were annotated to the gene with the nearest transcription start site using data tables downloaded from Ensembl BioMart (download as of 28.04.2020; Zerbino et al., 2018). Functional gene enrichment analysis was performed with genes containing differentially methylated CpGs or CpHs. For pathway analysis, we used the R package ReactomePA (Yu and He, 2016). SYNGO enrichment was performed using the online tool at https://www.syngoportal.org/. All enrichment analyses were performed with ENTREZ identifiers of differentially methylated genes as query lists and all genes covered by RRBS as background lists.

For transcription factor motif analysis, the number of differentially methylated CpGs that overlap with position weight matrices of transcription factor binding motifs was determined using the *Biostrings* package with a minimum match score of 90 %. Transcription factor motifs were retrieved from motifDb (Shannon and Richards, 2019). Motif enrichment for each transcription factor was tested by applying hypergeometric tests R. Multiple testing correction of *p*-values was performed using the FDR method if not stated otherwise.

#### Single-cell RNA sequencing data

Single-cell data were demultiplexed using bcl2fastq2 v2.20 and aligned to the mouse reference transcriptome mm10 using kallisto v0.44.0 with default parameters. Transcript levels were quantified as transcripts per million reads (TPM). TPM counts were imported into R and transcript information was summarized on gene-levels. We removed genes which did not have more than 10 reads for more than 500 genes. The pagoda2 package (Fan et al., 2016) was used for variance normalisation (gam.k = 10), identification of cell clusters (PCA: nPcs = 100, n.odgenes = 3e3; k-nearest neighbor: k = 50, perplexity = 30, min.group.size = 10) and *t-*distributed stochastic neighbor embedding (*t*-SNE). Clusters were annotated based on marker genes and two smaller clusters containing contaminating microglia and oligodendrocytes were removed and the clustering recalculated to yield 6 clusters covering the neurogenic trajectory. Log-transformed normalised gene expression values were then exported from pagoda2 for further analysis in R. Differential expression analysis was performed using edgeR (McCarthy et al., 2012) on the untransformed count data which had been adjusted using the zinbwave package (Risso et al., 2018).

Heatmaps were generated using custom code and data were either normalised to the T0 cluster or presented as KO/WT ratios as noted in the figure legends.

### Resource availability

Sequencing data will be deposited on GEO and made available upon publication. Code for bioinformatic analysis is available upon request.

### Statistics

Statistics was preformed using R v3.6.3 or GraphPad Prism v6. All experiments were performed with the experimenter being blinded to the experimental groups.

## Supplemental figures

**Fig. S1:**
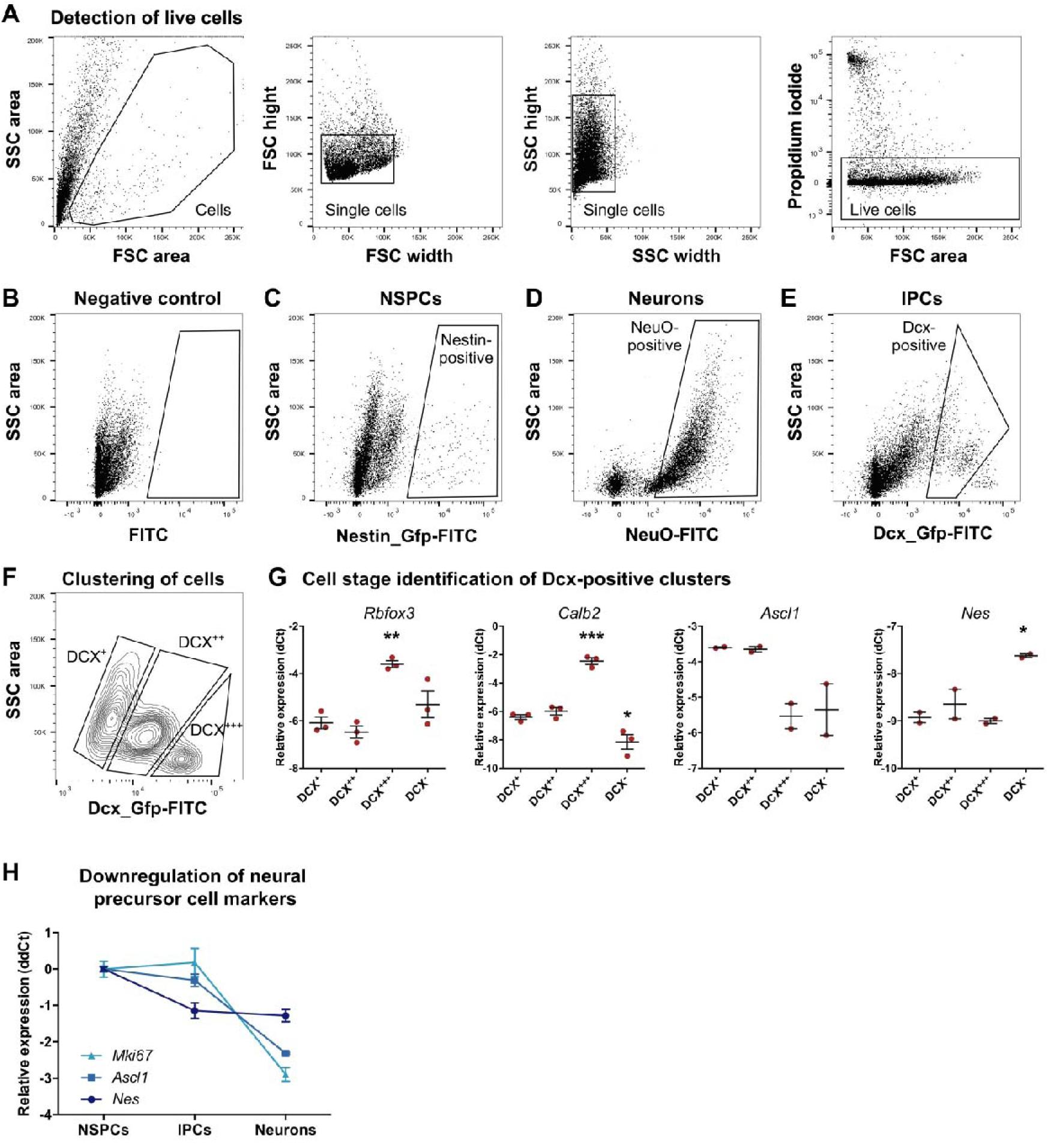
FACS strategy used for isolation of neural precursor cell populations and neurons from the adult dentate gyrus. **A,** Single cells were detected by plotting forward scatter (FSC) and side scatter (SSC). Dead cells were excluded based on incorporation of propidium iodide. **B,** Background Gfp intensity in a wildtype mouse. **C,** Neural stem and early progenitor cells (NSPCs) were isolated from *nestin*::GFP mice based on Gfp intensity. **D,** Neurons were identified based on incorporation of the neuron-specific dye NeuroFluor (NeuO). **E,** Late progenitor cells (lPCs) were identified as a sub-population of Gfp-positive cells isolated from *Dcx*::GFP mice. **F,** Dcx-GFP-positive cells cluster into three sub-populations that differ by GFP intensity and granularity (side scatter). **G,** Expression of marker genes relative to *Actb* in the different Dcx-positive cell clusters and Dcx-negative cells (DCX^-^) as determined by quantitative PCR. While the cluster DCX^+++^ showed higher expression of neuronal markers *Rbfox3* and *Calb2* compared to cluster DCX^+^, cluster DCX^++^ could not be distinguished from DCX^+^ based on the analyzed neuronal or precursor cell markers. DCX^+^ and DCX^++^ additionally showed higher expression of precursor cell marker *Ascl1*, identifying them as progenitor cells. Therefore, only DCX^+^ and DCX^++^ were isolated as lPCs as depicted in E. One-way ANOVA compared to DCX^+^ with *post hoc* Dunnett’s test (* *p* < 0.05; ** *p* < 0.01; *** *p* < 0.001). **H,** Downregulation of precursor cell markers *Nes* and *Ascl1* and proliferation marker *Mki67* during neuronal differentiation. Depicted are means with standard error of the mean.

**Fig. S2:**
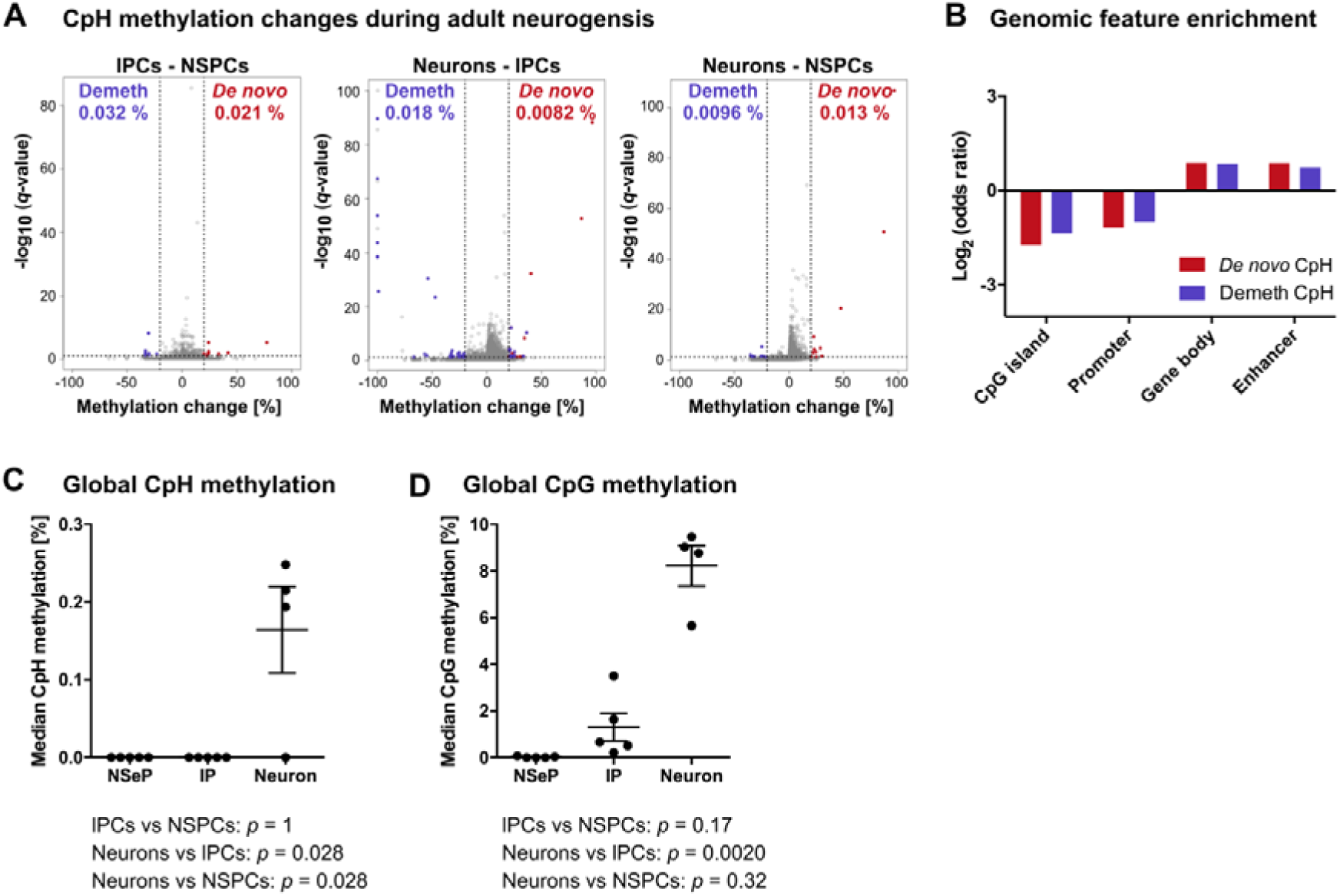
CpH methylation changes during adult hippocampal neurogenesis. **A,** Volcano plots depicting CpH methylation changes between NSPCs, lPCs and neurons of the adult dentate gyrus. Significantly differentially methylated cytosines with *q* < 0.05 and absolute methylation difference greater than 20 % are highlighted in violet (demethylated CpHs – Demeth) and red (*de novo* methylated CpHs - *De novo*). Percentage of differentially methylated CpHs among all sequenced CpHs are indicated in brackets. **B,** Differentially methylated CpHs were depleted at CpG islands and promoters and enriched at gene bodies (adjusted *p* < 0.05). No significant enrichment was found at enhancers (adjusted *p* > 0.05). **C,** The median CpH methylation levels over 1,849 CpHs covered by all samples was higher in neurons. **D,** Global methylation over 765 CpGs covered by all samples increased during differentiation. (*p*-values in C-D are from Kruskal-Wallis with *post hoc* Dunn’s test).

**Fig. S3:**
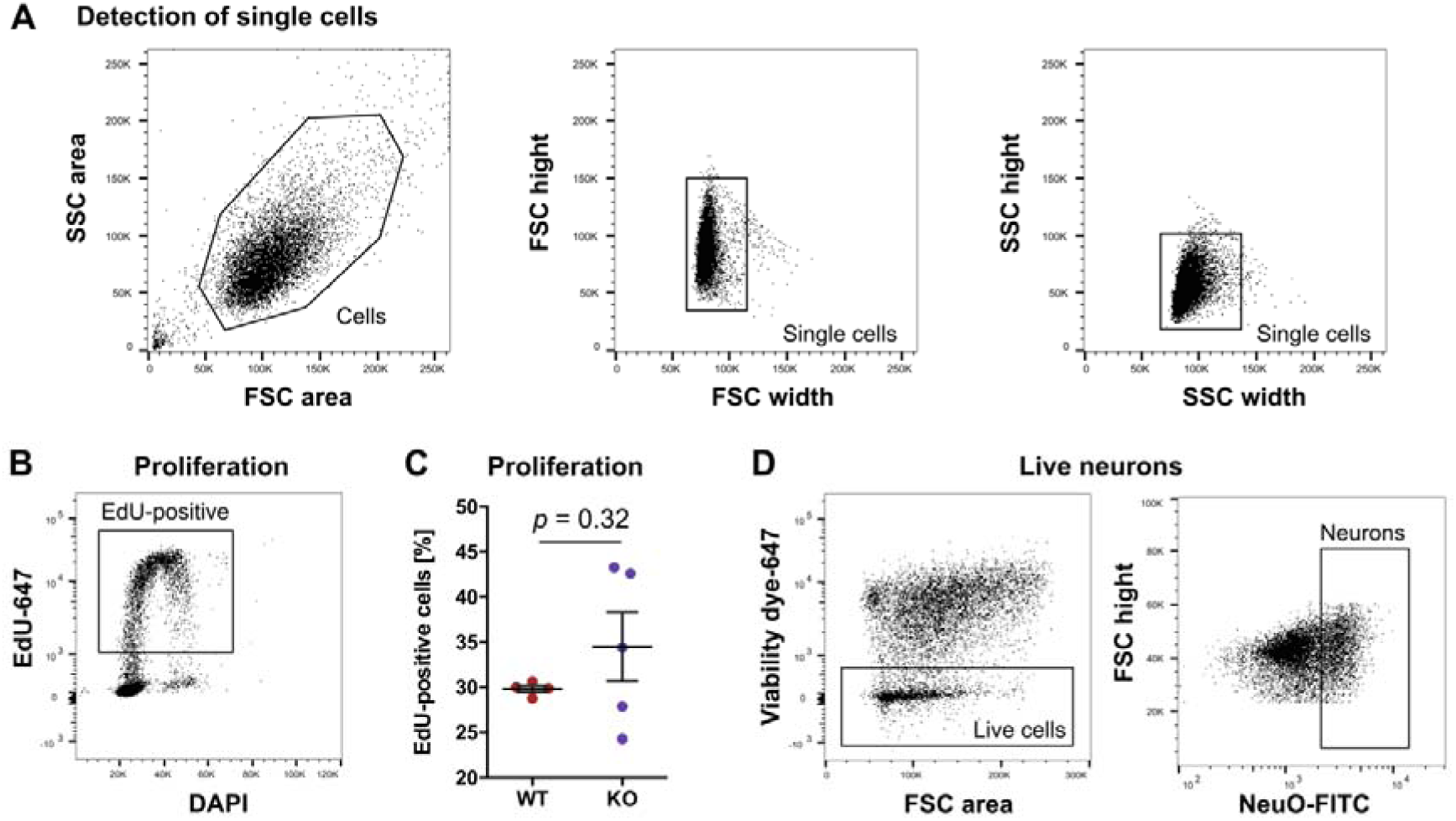
Analysis and isolation of *in vitro* neural precursor cells and differentiated neurons by flow cytometry. **A,** FACS strategy used for detection of neural precursor cells. Single cells were identified by plotting forward scatter (FSC) and side scatter (SSC). **B,** Cells in S-phase were identified based on EdU fluorescence intensity. **C,** Knock-out of *de novo* DNA methyltransferases does not affect neural precursor cell proliferation *in vitro*. **D,** Dead cells were removed from differentiated cultures using a viability dye and neurons were identified by fluorescence intensity of the neuronal dye NeuO in live cells (D corresponds to Fig. 3).

**Fig. S4:**
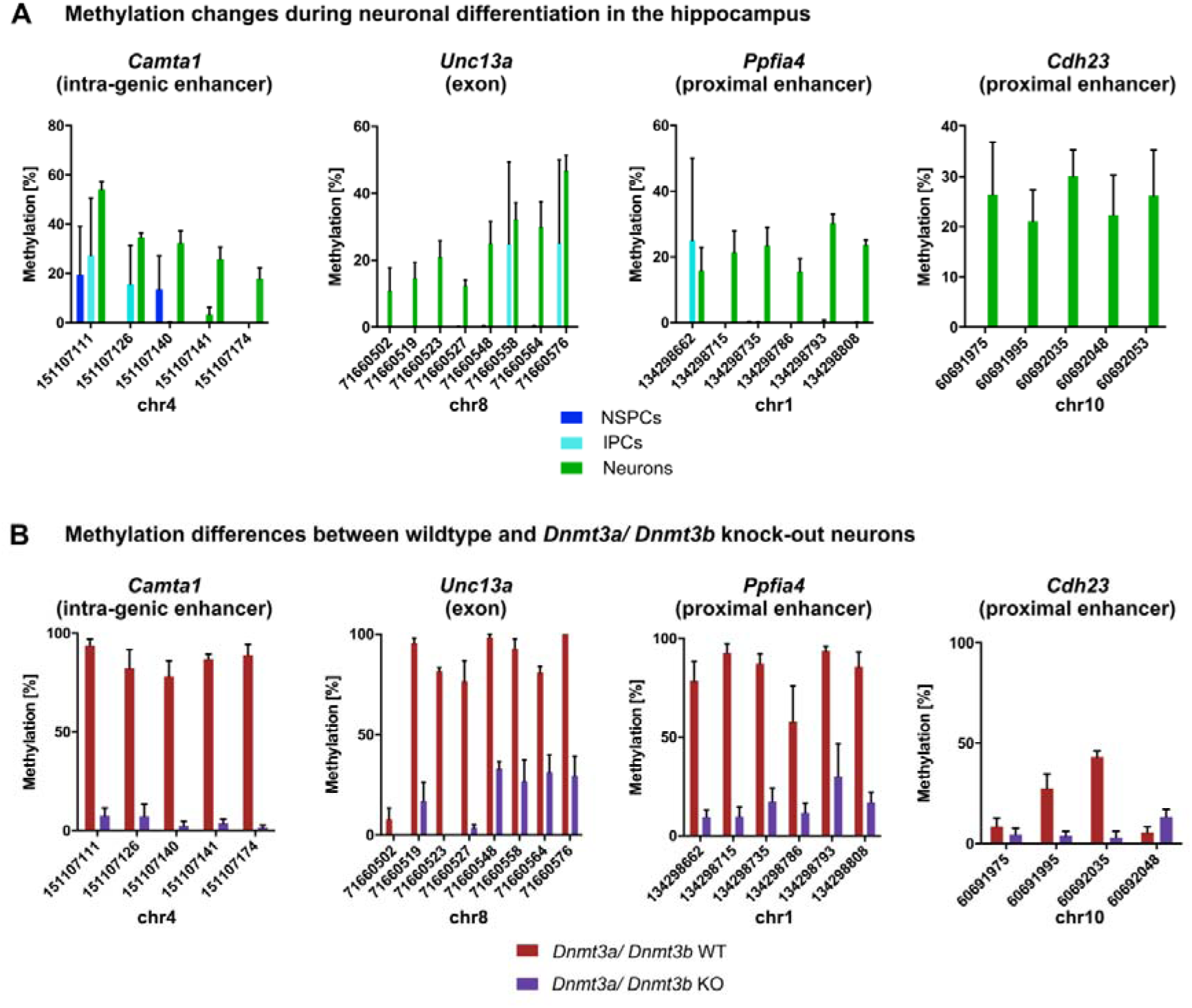
D*e novo* DNA methylation during adult hippocampal neurogenesis is mediated by *de novo* DNA methyltransferases. Depicted are representative genes that gain methylation during neuronal differentiation in the hippocampus (**A**) and are hypomethylated in neurons from *Dnmt3a/ Dnmt3b* KO neural precursor cells compared to WT neurons (**B**).

**Fig. S5:**
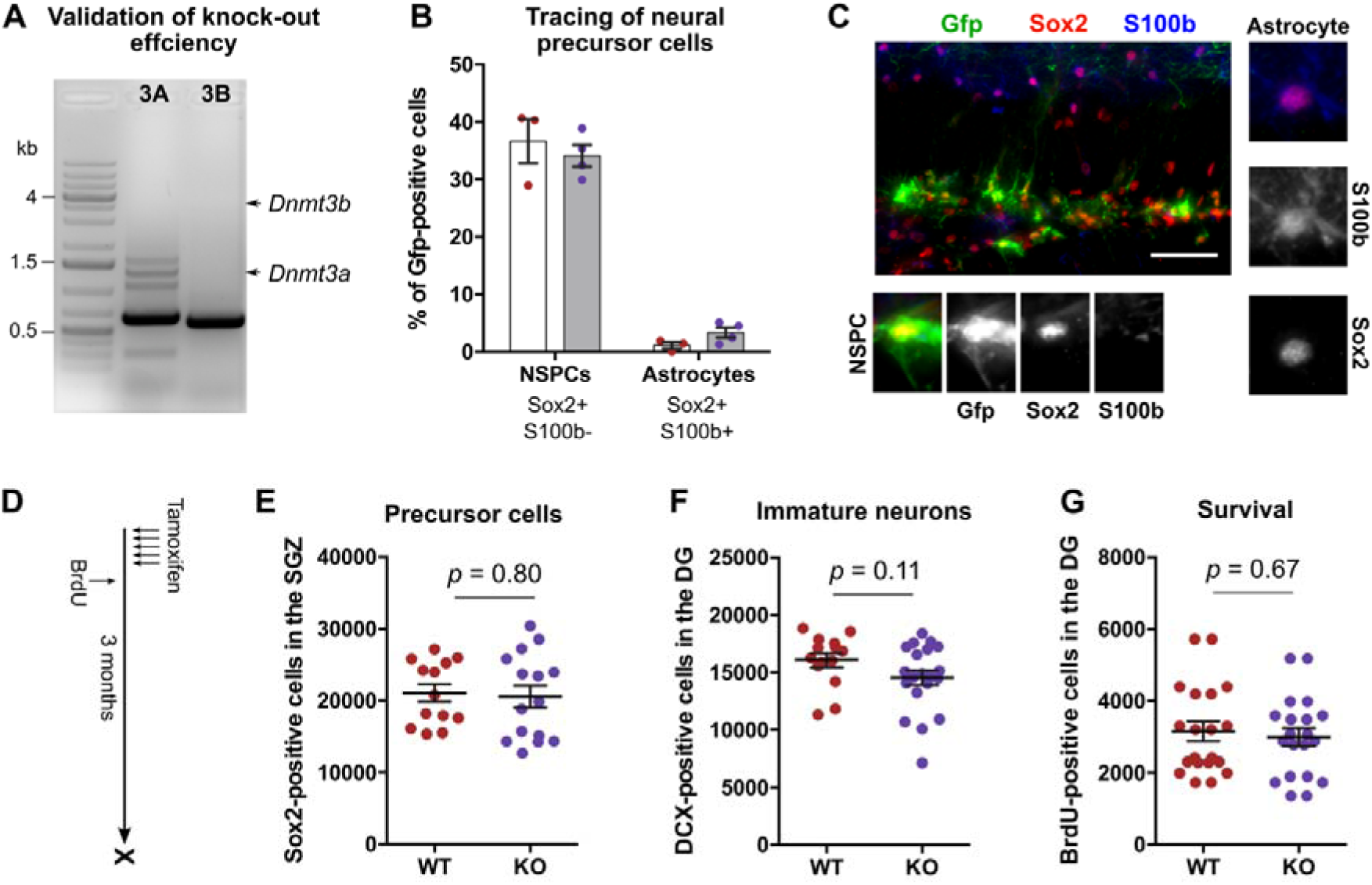
Deletion of *de novo* DNA methyltransferases does not influence long-term maintenance and potential of neural precursor cells in the hippocampus. **A,** Validation of deletion of *Dnmt3a* (3A) and *Dnmt3b* (3B) in Gfp-positive cells by polymerase chain reaction. Arrows indicate expected sizes for wildtype *Dnmt3a* and *Dnmt3b* amplicons. **B,** No difference in the percentage of NSPCs and astrocytes among Gfp-positive cells was detected between WT (red) and KO mice (violet). **C,** Representative fluorescent image for detection of NSPCs and astrocytes. Scale bar: 50 µm. D, Experimental protocol for results presented in E-G. No difference was found between WT and KO mice three months after tamoxifen administration in the total numbers of NSPCs in the sub-granular zone (SGZ; **E**), the numbers of new-born, immature neurons in the dentate gyrus (DG; **F**) or in the numbers of long-term survived cells (**G**).

**Fig. S6:**
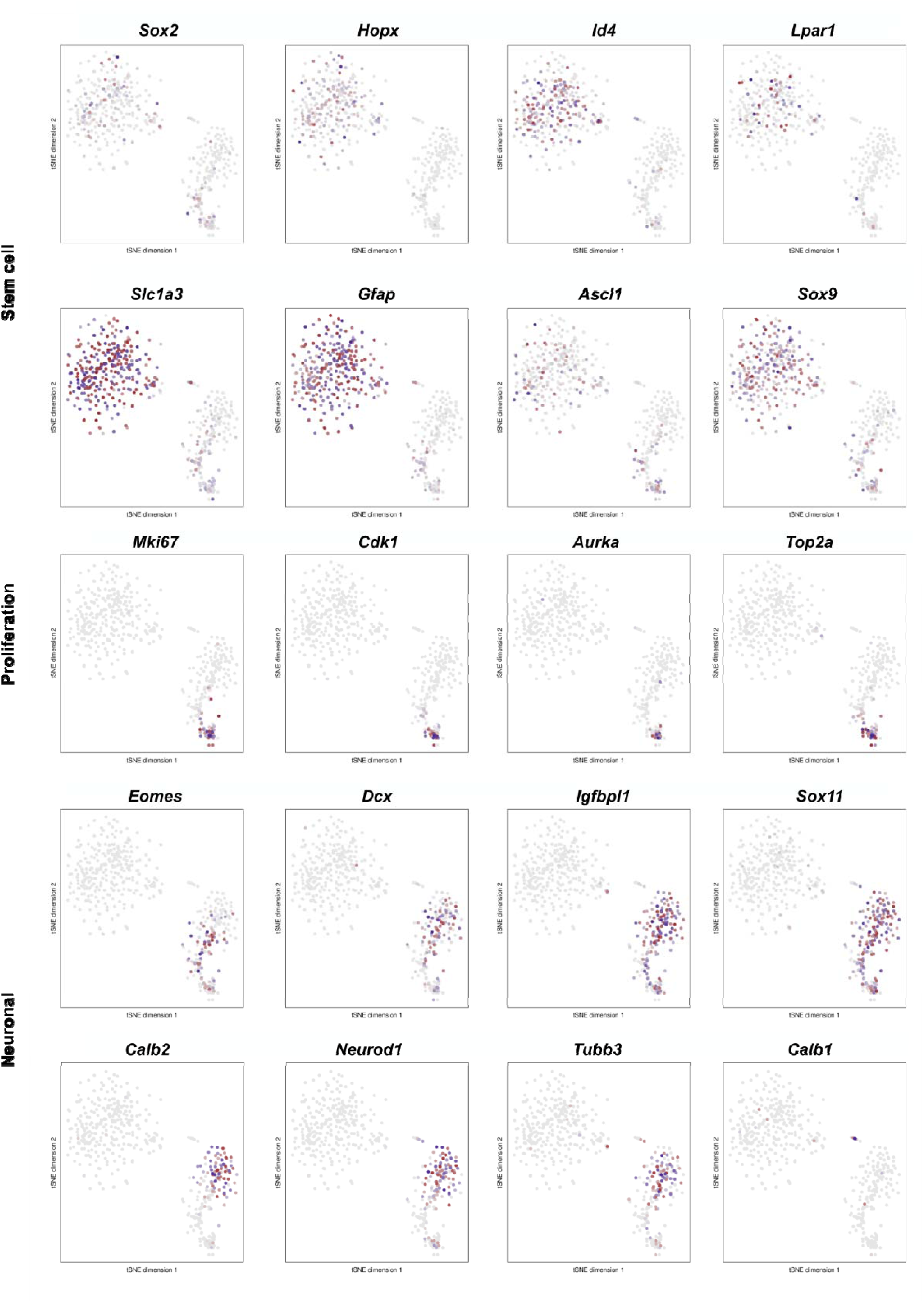
Marker gene expression in cell clusters. Depicted are t-SNE plots for marker genes that label specific cell stages during adult hippocampal neurogenesis. WT cells are highlighted in red, KO cells are labeled in violet.

**Fig. S7:**
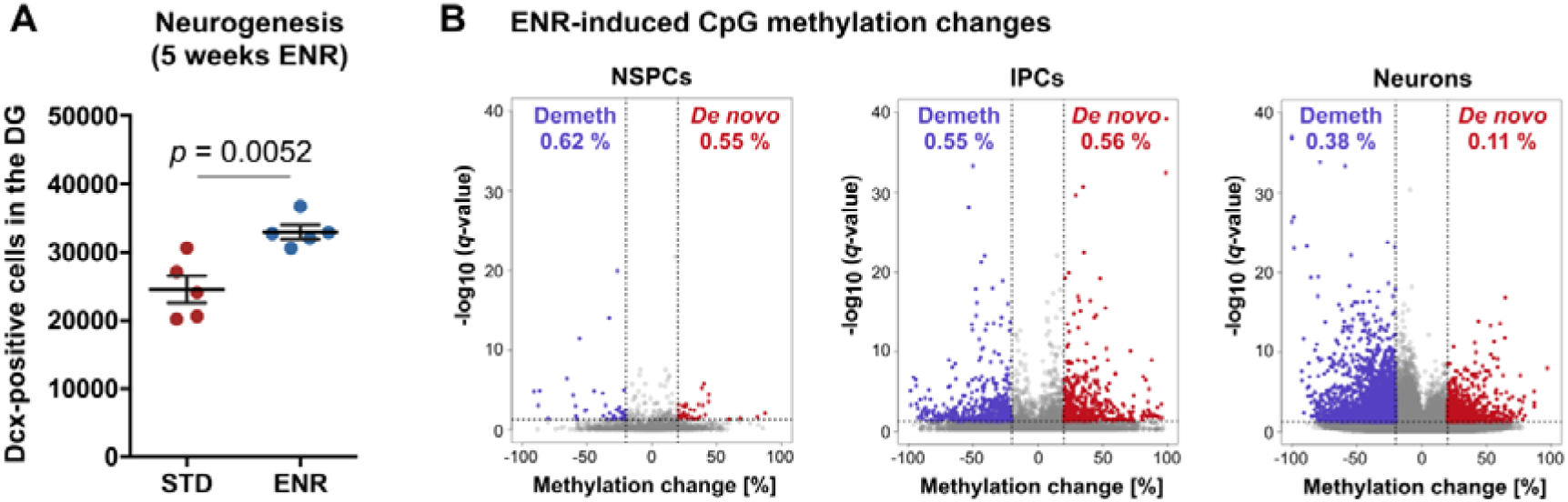
ENR-induced DNA methylation changes in hippocampal neural precursor cells and neurons. **A,** Increased numbers of immature (Dcx-positive) neurons in the dentate gyrus (DG) of C57BL/6JRj mice housed in environmental enrichment (ENR) for five weeks compared to mice housed in standard cages (STD). **B,** Differentially methylated CpGs in NSPCs, lPCs and neurons after five weeks of ENR housing. STD housed mice in B were identical to the group presented in Fig. 1.

**Fig. S8:**
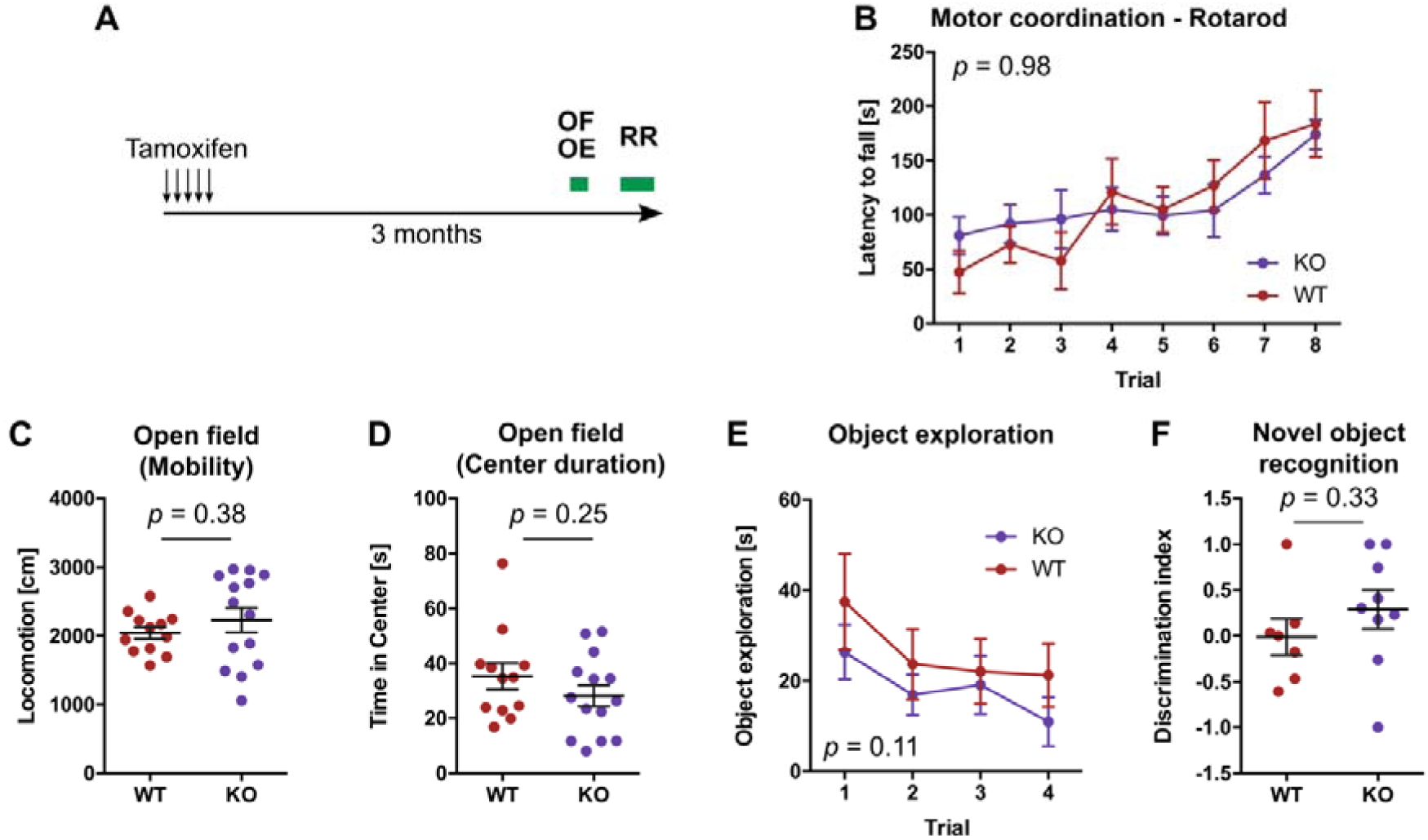
Deletion of *Dnmt3a* and *Dnmt3b* in adult neural stem cells and their progeny does not influence motor coordination, exploratory activity and recognition memory. **A,** Three months after administration of tamoxifen, mice were tested in one trial of open field test (OF) followed by four trails of object exploration test (OE). One week later, mice were tested on the rotarod (RR). **B,** Rotarod performance in WT and KO mice. Depicted *p*-value corresponds to genotype effect from repeated measure two-way ANOVA. **C,** Distance mice moved in the open field. **D,** Time mice spent in the center of the open field. **E,** Time mice spent around objects. In trial 4, one object was replaced with a novel object. Depicted *p*-value corresponds to genotype effect from two-way ANOVA. **F,** Relative time mice spent around the novel object in trial 4 (novel object - old object)/(novel object + old object). Depicted are means ± standard error of the mean. *P*-values in C, D and F are from unpaired *t*-test.

